# Circadian Clock Gene Modulation and Selective Reprogramming in Response to Virus Infection

**DOI:** 10.64898/2026.08.04.742878

**Authors:** Nicole Rivera-Espinal, Emna Achouri, Matthew Hackbart, Maria F. Gonzalez-Aponte, Yanling Yang, Erik D. Herzog, Carolina B. López

## Abstract

The circadian clock regulates fundamental cellular processes that include the host response to infection. However, the intersection of the circadian clock and the mechanisms that drive antiviral immunity remains poorly understood. Copy-back viral genomes (cbVGs) generated during virus replication strongly stimulate immunity during infection with negative-sense RNA viruses. Here, we demonstrate at the single cell level that A549 lung epithelial cells, commonly used to model lung infections, sustain circadian rhythms and upon infection with Sendai virus (SeV) progressively remodel the circadian clock through an innate immune response driven primarily by cbVGs. Furthermore, we show that cbVG-driven changes in circadian gene expression are mediated through two distinct innate immune sensing pathways: the RIG-I adaptor MAVS is required for preferential induction of the BMAL1 paralog *ARNTL2*, whereas the double-stranded RNA sensor PKR is required for the cbVG-specific induction of the negative-feedback regulators *NR1D1*, *NR1D2*, and *PER1*. A similar cbVG-specific signature was observed during respiratory syncytial virus infection, suggesting that cbVG-driven circadian gene regulation is not unique to SeV. In addition, we established through gain- and loss-of-function experiments that *ARNTL2* is functionally required for amplifying the transcriptional response to infection, with selective effects on the expression of specific antiviral genes, including *CCL5*. Together, these findings establish innate immune signaling triggered by cbVGs as selective driver of circadian clock gene expression during viral infection through distinct sensing pathways and identify ARNTL2 as a previously unrecognized regulator of virus-induced host transcriptional responses.

## INTRODUCTION

During viral infections, individual cells face distinct viral stimuli depending on the class and composition of viral RNA they encounter (1–4). In infections with non-segmented negative-sense RNA viruses, in addition to standard full-length viral genomes, the viral polymerase generates copy-back viral genomes (cbVGs) when it dissociates from the template and reinitiates replication using the nascent strand as a template (1, 2). These truncated and rearranged RNA molecules lack protein-coding capacity but can be packaged into virions and infect neighboring cells creating a heterogeneous population of infected cells with qualitatively distinct fates (5–8). cbVGs carry potent immunostimulatory motifs that are strong activators of cytosolic innate immune sensing pathways, most prominently the RIG-I/MAVS signaling pathway (5, 9–14), which drives antiviral responses, and the double-stranded RNA-activated kinase PKR (7), which mediates translational arrest, cellular stress responses, and broad transcriptional reprogramming. Through these pathways, cbVGs shape infection outcomes in distinct ways compared to standard viral genomes, making the viral RNA composition of the infected cell a key, and often overlooked, determinant of the host response (1, 6).

A host regulatory system relevant to infection outcomes is the circadian clock, a cell-autonomous molecular timekeeper that coordinates gene expression and physiology across approximately 24-hour cycles in all cells and tissues (15). At its core, this cell-intrinsic rhythmicity is maintained by a highly conserved transcriptional-translational feedback loop (TTFL). This loop is driven by factors including the heterodimeric activator complex CLOCK:BMAL1 (encoded by the genes *CLOCK* and *ARNTL1*), which bind to E-box elements in DNA to drive the rhythmic transcription of its own repressors, including the Period *(PER1/2/3*) and Cryptochrome (*CRY1/2*) families, as well as the nuclear receptors REV-ERB α/β (*NR1D1/2*) and ROR α/β/γ (NR1F1/2/3) **(Figure 1A)** (15, 16). Once translated, these negative regulators traffic back into the nucleus and inhibit the CLOCK:BMAL1 complex, completing a self-sustained autonomous cycle. Circadian-regulated genes are estimated to comprise roughly half of the mammalian transcriptome (17). Beyond timekeeping, this network directly coordinates many of the same pathways induced by cbVGs during replication, including innate immune signaling cascades (18–23), metabolic homeostatic pathways (24–26), translational control networks (27–30), and genome-wide transcriptional programming (17, 31–33).

**Figure 1.**
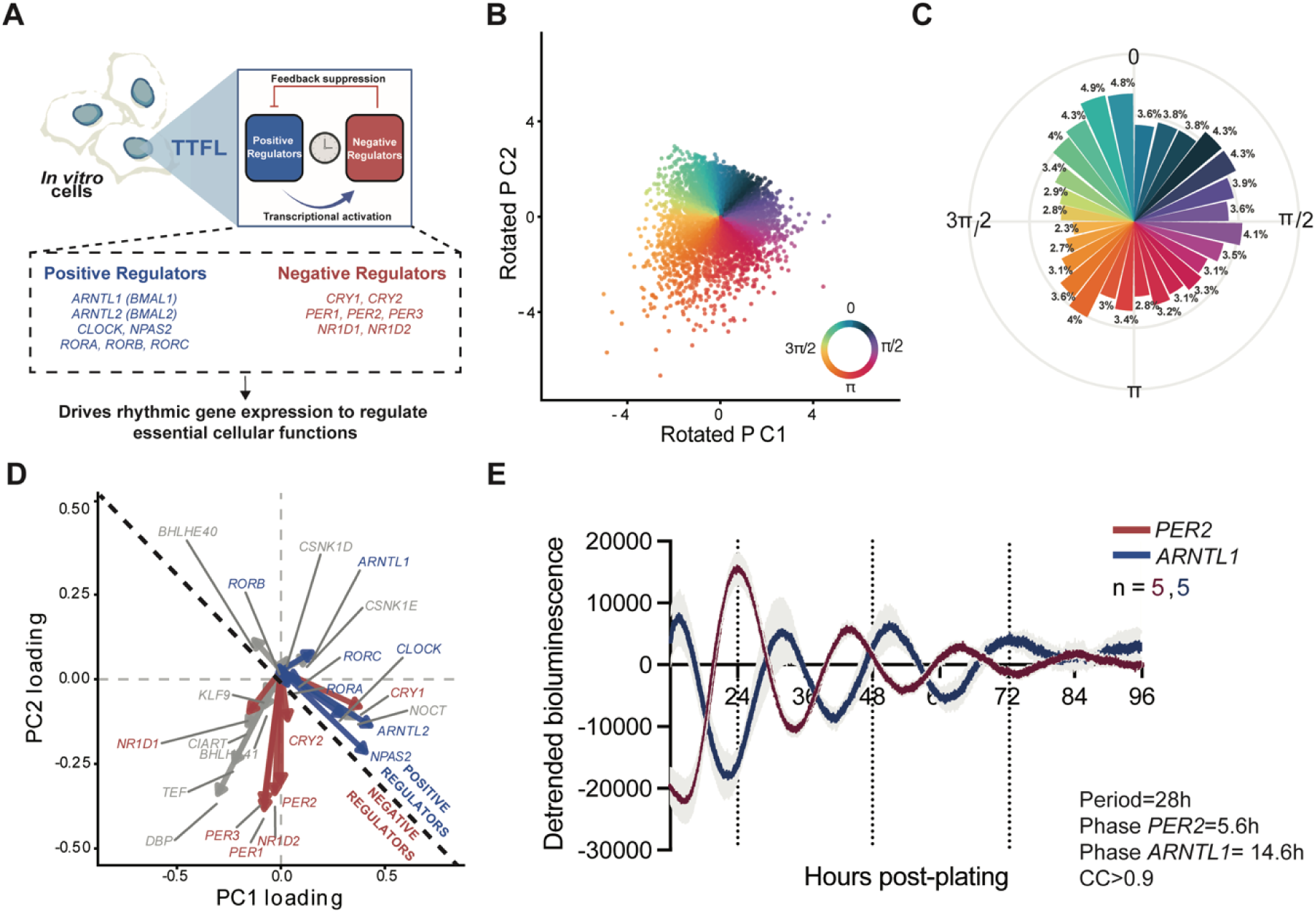
A549 cells retain a functional circadian clock. **(A)** Schematic of the transcriptional–translational feedback loop (TTFL), showing activating and repressive arms that drive rhythmic gene expression. **(B)** PCA projection of 5,000 mock-infected A549 cells onto a rotated PC1–PC2 coordinate space derived from 23 core circadian regulators. Each dot represents a single cell, colored by its inferred circadian phase. **(C)** Polar histogram showing the frequency of cells across the circadian phase (0–2π radians). The 2π radian circular space is divided into 28 equidistant bins. Bar height corresponds to absolute cell frequency, ranging between 117 and 244 cells per bin. Concentric grid lines indicate increments of 100, 200 and 300 cells. Relative percentages are annotated atop each bin, demonstrating a near-uniform distribution of cells throughout the reconstructed cycle. **(D)** PCA loadings of the 23 clock genes illustrating their contribution to the rotated PC1–PC2 space. Arrows indicate gene-specific loadings and are colored according to their role. Core positive regulators are shown in blue, red represents the core negative regulators, gray represents auxiliary regulators or output genes. Black dashed line indicates the inferred functional axis separating the activating (e.g., *ARNTL1*) and repressive (e.g., *PER2*) components of the TTFL. (**E)** Detrended bioluminescence values recorded from A549 cells stably expressing *PER2*- or *ARNTL1*-driven luciferase reporters, demonstrating sustained circadian oscillations in vitro. Traces represent mean ± SEM (n = 5 dishes per reporter). Period and phase values are indicated in figure; rhythmicity was assessed by cosine fitting (correlation coefficient, CC > 0.9).

Given this functional overlap, a growing body of literature suggests that genetic or pharmacological disruption of individual clock components significantly alters viral infection outcomes (34–41). However, these phenotypic effects remain highly variable and context dependent. For instance, while classical paradigms suggest that *ARNTL1* (*BMAL1*) expression acts in a protective manner as a regulator of antiviral immunity (42), during influenza A virus (34, 35) and respiratory syncytial virus (RSV) (36), it exerts opposing effects on other viruses by restricting or altering the lifecycles of members of the *Flaviviridae* family, including hepatitis C, dengue, and Zika viruses (37). Similarly, negative regulators of the clock can exert context-dependent antiviral activity, with their manipulation selectively influencing viral replication depending on which clock regulator is being studied, the viral family, and cellular context (39, 43, 44). This profound variability suggests that circadian players do not exert a uniform antiviral function but instead regulate host responses in a virus- and context-dependent manner.

Despite well-established appreciation for the immunomodulatory capacities of copy-back viral genomes (cbVGs) and the increasing appreciation of the circadian clock in coordinating cell defense, these two fields have remained separate. Most studies examine circadian-virus interactions by treating infection as a homogeneous stimulus and the clock as a single, static functional unit, omitting how distinct viral RNA sub-species might selectively engage individual transcriptional-translational feedback loop (TTFL) components. This represents a critical gap in knowledge as the cytosolic innate immune pathways activated by cbVGs intersect directly with downstream transcriptional networks governing clock-regulated genes (30, 45–48). We hypothesize that the innate immune response engaged by distinct viral genome populations shapes circadian gene expression and activity during infection. Here, we track the molecular clock at single-cell resolution across an active infection timeline, profiling host transcriptomes sorted by specific intracellular viral genome architectures and functionally validating the distinct roles of copy-back and full-length viral species in remodeling individual clock components. By evaluating the molecular clock through these complementary layers, this work provides a direct look at how viral population heterogeneity actively reshapes individual components of the host timekeeping machinery, while defining the precise contributions of individual components in regulating host defense.

## RESULTS

### A549 lung epithelial cells maintain a functional circadian clock under infection-relevant culture conditions

Cell-intrinsic circadian gene oscillations have been reported to persist *in vitro*, where individual cells maintain rhythmic TTFL activity even without tissue-level synchrony (49). To determine how viral infection remodels circadian gene expression, however, measuring the average rhythm of an entire culture is insufficient because infected cells differ substantially in viral burden, viral genome composition, replication status, and innate immune activation. We therefore sought to use gene expression to infer the circadian phase of individual cells by developing a single-cell analytical framework that could be applied to data collected before and after infection. To do this, we adapted a principal component analysis (PCA)-based approach previously used to reconstruct cyclic transcriptional programs from single-cell RNA sequencing data (50–53). In this framework, cells progressing through a cyclic transcriptional program arrange along a continuous trajectory in a low-dimensional space. Depending on the underlying transcriptional variation, these trajectories can appear as arcs, ellipses, or circles, but all represent continuous progression through a periodic biological process (50–53). We first validated this approach using data from single-cell sequenced mock-infected A549 cells(54)and analyzing cell-cycle genes, which reconstructed the expected trajectory corresponding to canonical cell-cycle progression (**Supplemental Figure 1A**).

Having established that this approach accurately reconstructed a known cyclic transcriptional program, we next applied it to circadian gene expression. We analyzed a random subset of 5,000 A549 cells that were mock infected by changing their media 24 h after plating and collected for analysis 6h after mock infection (i.e. 30 h after plating and 6h after a media change). Single-cell RNA data was obtained from a previously published dataset (54). To reconstruct circadian organization, we used a curated panel of 23 genes spanning the major components of the transcriptional-translational feedback loop, including positive regulators (*ARNTL1*, *ARNTL2*, *CLOCK*, *NPAS2*, and *RORA-C*), negative regulators (*PER1-3*, *CRY1-2*, *NR1D1-2*), auxiliary regulators (*CIART*, *BHLHE40-41*, and *KLF9*), and clock-controlled output genes (*DBP*, *TEF*, *NOCT*, and *CSNK1D-E*) (**Supplemental Figure 1B**; see Methods for the complete gene panel and selection criteria). The resulting projection organized cells along a continuous circular trajectory in PC1-PC2 space (**Figure 1B**), consistent with a cyclic transcriptional program. To facilitate comparisons across experimental conditions, we established a common phase reference by orienting the reconstructed trajectory so that θ = π corresponded to the inferred peak of *PER2* expression in each cell, a well-characterized E-box target that was reliably detected in the single-cell dataset (55). To visualize the distribution of cells along the reconstructed trajectory, the circular projection was partitioned into 28 equal phase intervals (**Figure 1C**). The 5,000 mock-treated cells used for trajectory reconstruction were distributed across all phase intervals, with most containing approximately 3–4% of the total population.

We then examined whether the distribution of circadian genes within the PC1-PC2 space was consistent with the established organization of the clock gene expression (**Figure 1D**). In this representation, vector direction indicates the relative phase of the inferred daily maximal gene expression, whereas vector length reflects each gene’s contribution to the reconstructed transcriptional program. Genes with related functions clustered together within the reconstructed space, consistent with their known temporal relationship within the circadian cycle (56). CRY1 was the notable exception, clustering closer to genes within the positive regulatory arm of the clock, than to the other negative regulators, consistent with its previously described delayed transcriptional dynamics and complex regulatory architecture (57, 58). Together, these relationships indicate that the reconstructed trajectory preserves the expected transcriptional organization of the circadian clock in single A549 cells.

To test if the inferred phase order of uninfected cells aligns with their circadian rhythmicity, we next recorded bioluminescent reporters of two canonical circadian clock genes (*ARNTL1*- or *PER2*-luciferase) from cultured A549 cells. Cells were monitored every 3 minutes for 96 hours following plating under the same infection-compatible culture conditions used for the single-cell experiments (**Figure 1E and Supplemental Figure 1C**). Both reporters exhibited sustained circadian oscillations with strong rhythmicity (CC > 0.9) and an average period of approximately 28 hours. *PER2* and *ARNTL1* reached peak expression 5.6 and 14.6 hours after plating, respectively, corresponding to an approximately 9-hour phase difference, consistent with their relative positioning in the reconstructed trajectory (**Figure 1D**). Importantly, these population-level daily oscillations dampened over time, consistent with our finding that single cells had different circadian phases at 30 hours after plating when the single-cell RNA sequencing was performed (**Supplemental Figure 1C)**, when population-level oscillations remained robust but had begun to dampen (**Figure 1E**). Thus, while the culture retained measurable circadian rhythmicity, the increased variation in phase across individual cells likely enabled reconstruction of the circadian transcriptional continuum from a single-cell snapshot. These observations are consistent with previous work showing that A549 cells possess a functional circadian clock capable of driving rhythmic gene expression (59) supporting the use of this framework to infer the relative circadian organization of individual cells throughout infection.

### Sendai (SeV) infection induces transient transcriptional reorganization and progressive spatial distortion of the circadian clock

Having established a framework to infer the circadian clock organization of individual A549 cells, we next asked how SeV infection remodels this organization over the course of infection. We applied the circadian phase inference framework to SeV-infected cells from our previously published single-cell RNA sequencing dataset (54), and first compared the oscillatory behavior of individual circadian clock genes between mock- and 24 hpi, the latest time point available (**Supplemental Figure 1B)**. Gene expression along the inferred circadian trajectory was fit to a sinusoidal model and quantified using cosine correlation coefficients (CC), with a CC ≥ 0.70 defined as circadian.

We found oscillatory disruptions manifested as a reduction in sinusoidal coherence, a shift in peak expression phase, and a change in amplitude across multiple circadian clock components at 24 hpi. Among activating components, *CLOCK* showed the most pronounced loss of coherence, with its CC dropping well below the rhythmicity threshold (CC: 0.94 to 0.32). *ARNTL1* (CC: 0.84 to 0.66) and *NPAS2* (CC: 0.97 to 0.68) also showed reduced coherence, falling just below the threshold by 24 hpi (**Figure 2A**; **Supplemental Figure 2A**). *ARNTL2* was a striking exception, maintaining near-perfect sinusoidal coherence throughout infection (CC: 0.98 to 0.97), while exhibiting a clear directional shift in its peak expression phase relative to mock. Repressive components largely retained their coherence above the threshold, showing only modest reductions across *PER1* (CC: 0.91 to 0.77), *PER2* (CC: 0.96 to 0.94), *PER3* (CC: 0.91 to 0.84), and *NR1D1* (CC: 0.87 → 0.79). *CRY1* displayed a distinct phase shift, while *NOCT* exhibited a complete inversion of its expression pattern relative to mock, similar to *ARNTL2* (**Supplemental Figure 2B**). Overall, these findings indicate that SeV infection does not uniformly disrupt the circadian clock, but instead selectively rewires its transcriptional program relative to basal conditions, altering both the timing and strength of oscillatory expression across individual clock components.

**Figure 2.**
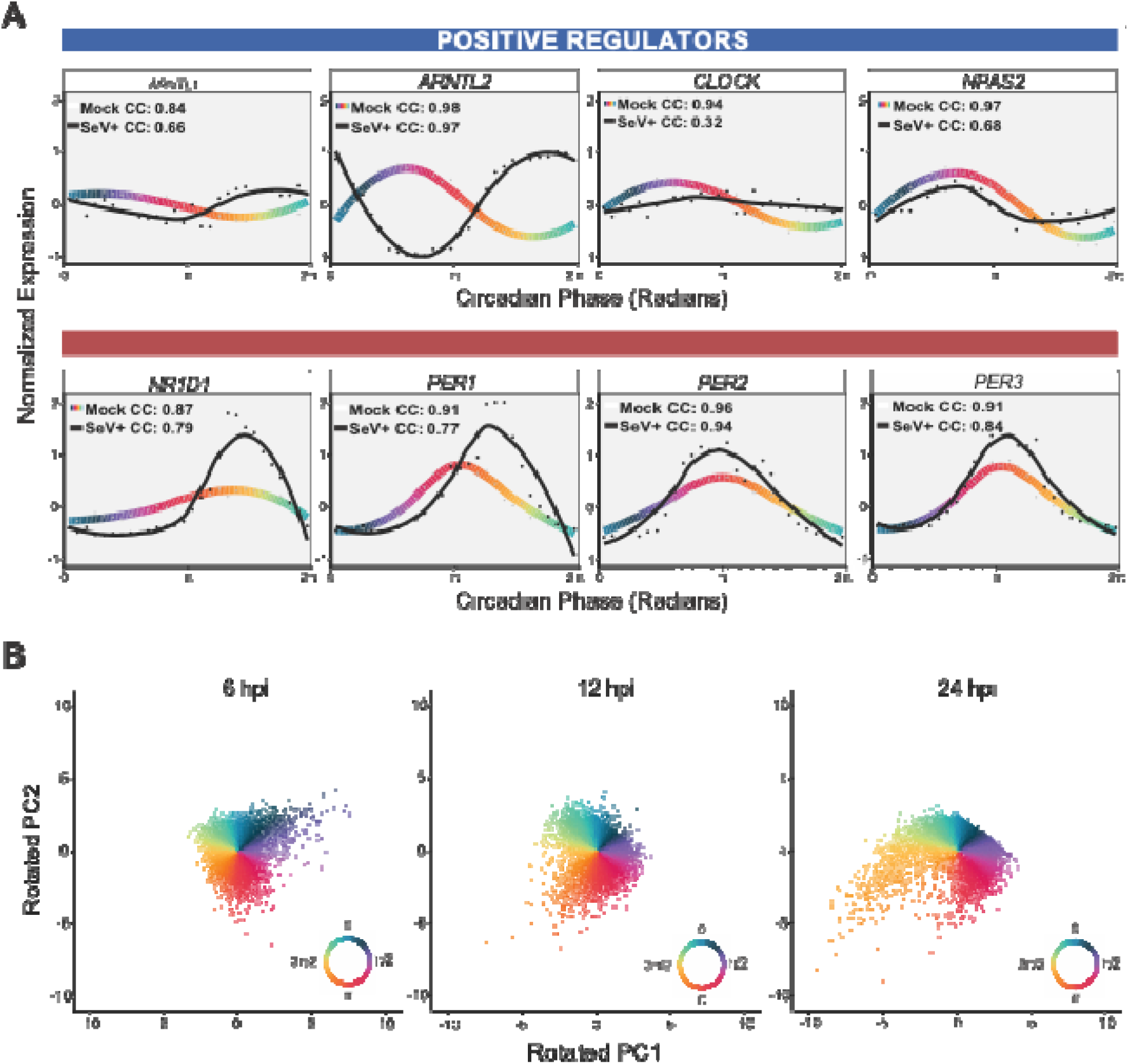
Expression profiles of positive and negative circadian clock regulators along the inferred circadian trajectory. **(A)** Average normalized expression of representative positive (blue) and negative (blue) circadian clock components across 28 equidistant phase bins. Individual dots represent the mean expression of each bin (n=179 cell/bin) for mock (gray) or 24hpi (black) conditions. Solid lines represent a LOESS smoothing (span = 0.7) of binned mean expression values, with line color indicating the inferred circadian phase (0–2π) of the mock cells. Smoothed expression curves are shown in black for the 24hpi timepoint. Cosine correlation coefficients (CC) indicate goodness of sinusoidal fit for each gene. **B)** PCA projection maps of single-cell host transcriptomes at 6, 12, and 24 hpi. Cells from each timepoint are projected onto the rotated PC1-PC2 coordinate space established by the 23-gene clock profile and colored by their position along the circadian phase vector (inset color wheels), tracking progressive clock disruption across the infection timeline.

To determine whether the transcriptional changes observed at 24 hpi progressed throughout the infection course, we next examined the reconstructed circadian organization across the SeV infection time course (**Figure 2B**). These timepoints correspond to the period during which the luciferase recordings demonstrated sustained anti-phase *PER2* and *ARNTL1* oscillations, providing a circadian reference for interpreting changes in the reconstructed single-cell trajectories. Relative to mock cells (**Figure 1D**), the position of circadian gene vectors, representing the inferred phase of maximal gene expression, changed throughout infection (**Supplemental Figure 2C**). These changes were evident by 6 hpi, reached their greatest deviation at 12 hpi, and partially returned toward the mock configuration by 24 hpi. In contrast, the distribution of cells across the inferred circadian trajectory changed progressively over time (**Figure 2B**). Cell enrichment within the π–3π/2 interval (represented by the orange-colored cell population on the PCA plot) increased from 6 to 24 hpi, producing a progressively elongated trajectory that was most pronounced at 24 hpi.

Together, these findings indicate that SeV infection selectively remodels circadian gene expression rather than uniformly disrupting it. Genes associated with the positive regulatory arm of the clock exhibited the greatest loss of oscillatory coherence, whereas genes associated with the negative regulatory arm largely retained rhythmicity. *ARNTL2* was a notable exception, maintaining strong rhythmic coherence despite a pronounced shift in peak expression phase. While changes in the inferred phase of maximal gene expression were most pronounced at 12 hpi, redistribution of cells across the reconstructed circadian trajectory progressed throughout infection, resulting in increasing enrichment within the π–3π/2 region of circadian pseudotime by 24 hpi.

### A distinct transcriptional circadian clock state associates with antiviral gene expression patterns and cbVG detection

To determine whether the circadian reorganization observed through our clock-specific PCA framework was due to global transcriptional changes, we applied UMAP dimensionality reduction to the complete single-cell RNA-seq dataset and projected the previously inferred circadian phase of each cell onto the global transcriptional UMAP (**Figure 3**). In mock-treated cells, cells spanning the full range of inferred circadian phases were broadly distributed throughout the global transcriptional landscape, with no preferential enrichment of any circadian phase within a particular transcriptional state (**Figure 3A**). In comparison, infected cells showed an expression distribution that became increasingly non-uniform as infection progressed (**Figure 3A**). At 6 hpi, a small population of cells began to separate in UMAP space showing a distinct enrichment for early circadian phases (0–π/2; dark blue and purple), while the remaining cells continued to span the rest of the inferred circadian cycle. By 12 hpi, this phase-enriched population expanded and became concentrated within the 3π/2–0 interval, whereas the remaining cells continued to occupy later circadian phases. By 24 hpi, this segregation became more pronounced, with discrete transcriptional regions exhibiting increasingly restricted circadian phase distributions that ranged from blue/green (0) to yellow (3π/2). Together, these observations indicate that infection progressively reorganizes the relationship between circadian phase and the global transcriptional landscape, resulting in preferential enrichment of specific circadian phases within distinct transcriptional populations.

**Figure 3.**
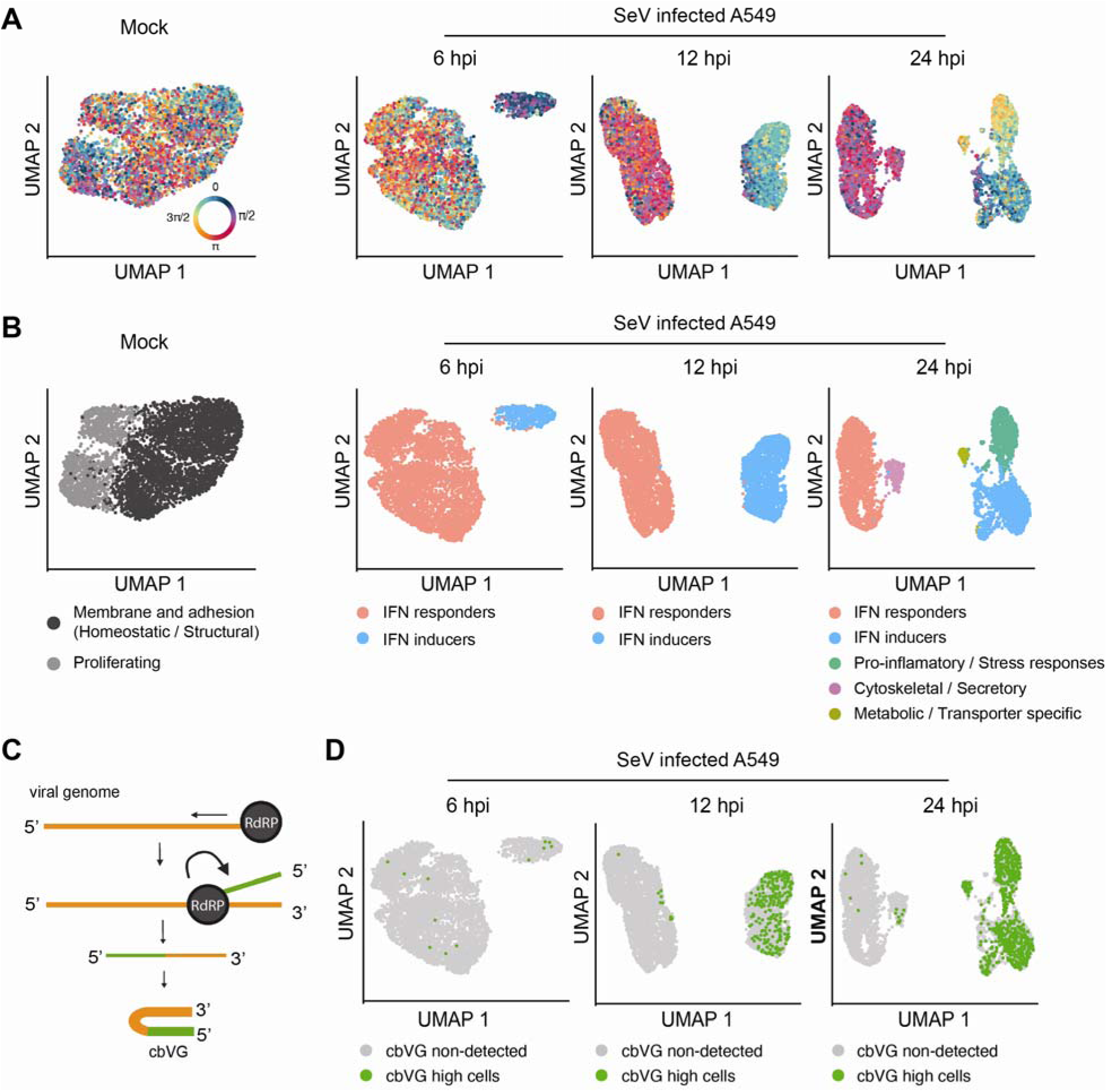
Single-cell mapping of circadian clock phase, transcriptional clusters, and cbVG accumulation during SeV infection. **(A)** UMAP projection of mock and SeV-infected A549 cells at 6, 12, and 24 hpi colored by the inferred circadian phase from the clock PCA framework. **(B)** UMAP projections colored by transcriptional cluster identity. Clusters were identified using the Louvain algorithm (Seurat FindClusters) at a resolution of 0.1 and annotated based on top 10 differentially expressed marker genes. **(C)** Schematic of copy-back viral genome (cbVG) formation during RNA virus replication. cbVGs arise when the viral RNA-dependent RNA polymerase (RdRP) reattaches to the nascent RNA strand, generating truncated self-complementary genomes. **(D)** UMAP projections of SeV-infected A549 cells at 6, 12, and 24 hpi with cbVG-high cells highlighted in green and cells without detected cbVG signal shown in gray.

To define the functional identities associated with these configurations, we employed the Louvain algorithm with a low-resolution parameter (res = 0.1) to identify the most robust, high-level cellular states (**Figure 3B**). To ensure clean and definitive functional annotations, clusters were defined based on their top differentially expressed markers and mapped onto a UMAP (**Supplemental Table S1-4 and Supplemental Figure 3**). As we previously reported (54), in mock cells, clusters corresponded to homeostatic populations, including membrane/adhesion (homeostatic/structural) and proliferating states. During infection, two dominant host-defense populations emerged: IFN inducers, characterized by the expression of type I and III interferon genes, and IFN responders, which were heavily enriched for downstream interferon-stimulated genes (ISGs) (54). By 24 hpi, host-cell transcription further diversified to include pro-inflammatory/stress, neuroendocrine-like (cytoskeletal/secretory), and metabolic/divergent transporter-specific populations. Overlaying our clock data onto these functional clusters revealed a striking, non-random alignment. Cells within the 3π/2 to 0 phase interval localized predominantly within the IFN-inducer population, whereas cells positioned in the π to π/2 interval mapped preferentially to IFN-responder states. At 24 hpi, the newly emerged pro-inflammatory and metabolic clusters packed tightly into the 3π/2 phase region, closely mirroring the spatial clustering of the core repressors *NR1D1* and *NR1D2* observed in our phase trajectory map (**Supplemental Figure 2C**). Together, these findings demonstrate that infection-induced shifts in the molecular clock correspond tightly to the activation of global immune and metabolic programs.

Because changes in the cell cycle can drastically alter the global host transcriptome (52), we examined whether the observed phase clustering reflected shifts in cell division rather than true clock remodeling (**Supplemental Figure 3)**. Mapping cell-cycle states onto infected cells revealed relatively stable distributions during early infection (6 -12 hpi), even as the clock phase enrichment became distinct and the cbVG-rich IFN-inducer population emerged. More pronounced shifts in cell-cycle composition, specifically an alteration in G1 and S/G2M proportions, became evident only later at 24 hpi (**Supplemental Figure 3)**. Because the positioning of cells into specific clock intervals and the appearance of IFN inducers happen prior to these bulk changes in cell division, cell-cycle progression is unlikely to account for the initial skewed clock phase observed during early infection.

The emergence of a distinct IFN-inducer population occupying a restricted region of the circadian pseudotime prompted us to ask whether these cells were enriched for cbVGs. cbVGs are non-standard viral genomes generated during RNA virus replication that act as potent immunostimulatory molecules and are the primary drivers of interferon production during SeV infection (1) (**Figure 3C**). Using a specialized library preparation strategy that enables the sequencing and computational analysis of cbVG sequences at single-cell resolution (54, 60), we identified cbVG-high cells and mapped their distribution in the UMAP space (**Figure 3D**). As previously reported (54), 6 hpi, cbVG-high cells were rare and mostly localized to peripheral regions of the UMAP space. As the infection progressed to 12 hpi, the population of cbVG-high cells concentrated within the IFN-inducer cluster. At 24 hpi, cbVG-high cells expanded beyond the IFN-inducer population, accumulating in the newly emerged pro-inflammatory and stress-associated clusters. As anticipated, not all cells within the IFN-inducer cluster were identified as cbVG-high, a reflection of the inherent sensitivity limitations of single-cell cbVG detection. Nevertheless, the detected cbVG-high cells consistently localized to a restricted region of transcriptional space that overlaps with the circadian clock configuration enriched near the 3π/2 to 0 interval. Together, these observations demonstrate a precise, tri-directional alignment between host circadian clock state, global antiviral immune activation, and intracellular viral genome configuration.

### cbVG accumulation selectively induces *Arntl2* and repressive clock gene expression components

To assess the relationship between viral genome composition and clock gene expression, we next analyzed a previously published bulk RNA-seq dataset from RNA FISH-IFA-sorted SeV-infected A549 cells, classified as cbVG-high, standard (std) genome-high, or non-detected (ND) (5) (**Supplemental Figure 4A**). This approach enriches for cells that have accumulated cbVGs and enables comparison of transcriptional programs associated with distinct viral genome species. Analysis of circadian gene expression across these populations revealed a distinct pattern in cbVG-high cells (**Supplemental Figure 4B**). Within this group, core repressive clock components were preferentially and robustly upregulated, including the negative feedback elements *PER1*, *NR1D1*, and *NR1D2*. This coordinated induction also extended to the auxiliary regulators *BHLHE41* and *KLF9* (**Supplemental Figure 4B**); notably, while categorized as auxiliary regulators rather than core clock components, both factors are well established to exert potent repressive roles on clock-driven transcription (61–63). In contrast to this broad induction of repressors, positive regulators of the core clock loop were largely unaffected, with *ARNTL2* representing the only activator that was significantly induced. Furthermore, the auxiliary clock-controlled output gene *NOCT* was identified as the most strongly upregulated transcript among all evaluated clock genes (**Supplemental Figure 4B**). This signature was highly specific to cells enriched in cbVGs, as cells characterized by high levels of standard genomes exhibited only modest transcriptional changes across a few positive regulators. Taken together, these data suggests that the accumulation of immunostimulatory cbVGs selectively drives a repressive clock state, characterized by the preferential upregulation of core transcriptional loop repressors alongside the robust, non-canonical induction of the *ARNTL2* activator paralog.

To validate whether cbVGs are sufficient and necessary to upregulate this clock transcriptional program, we infected A549 cells with SeV stocks lacking cbVGs (cbVG−) and compared them to cells infected with the same stocks supplemented with purified cbVG-containing particles, (cbVG+), as done previously (7, 64) (**Figure 4A**). Consistent with their established role, cbVG+ infection resulted in reduced expression of SeV nucleoprotein (NP) (**Figure 4B**) and markedly higher expression of the antiviral cytokine IL-29 (IFN-λ) relative to cbVG− infection (**Figure 4B**). Confirming that cbVGs potently activate innate immune signaling independently of productive viral replication and providing a robust system to confirm cbVG-dependent regulation of circadian clock gene expression during infection.

**Figure 4.**
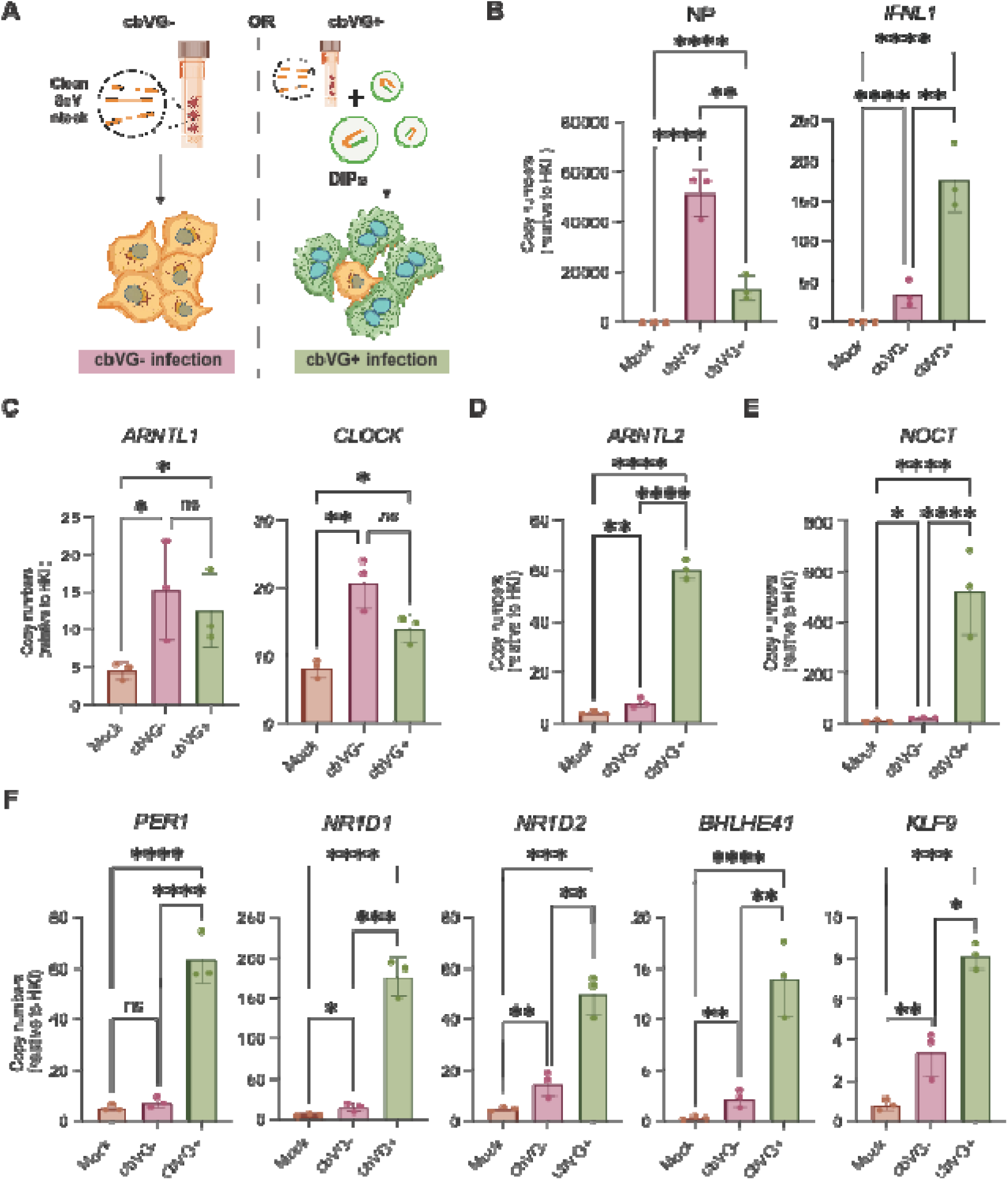
cbVGs preferentially induce repressive circadian clock components while selectively upregulating the activating gene *ARNTL2* and the clock-controlled output *NOCT*. **(A)** Schematic of experimental design. A549 cells were infected with SeV at an MOI of 1.5 TCID₅₀/cell using either a cbVG-negative (cbVG−) stock or the same stock supplemented with purified cbVG-containing particles “DIPs” (cbVG+) (purified cbVG particles at 20 HAUs). **(B)** mRNA expression of SeV nucleoprotein (NP) and IL-29 in A549 cells 24 h post infection (hpi) under cbVG− and cbVG+ infection conditions. **(C–F)** mRNA copy numbers of circadian clock genes in A549 cells at 24 hpi under mock, cbVG−, and cbVG+ conditions. Genes shown include core activating components (*ARNTL1*, *CLOCK*), the *ARNTL1* paralog *ARNTL2*, the clock-controlled output gene *NOCT*, and repressive/auxiliary regulators (*PER1*, *NR1D1*, *NR1D2*, *BHLHE41*, and *KLF9*). mRNA levels were normalized to housekeeping genes index “HKI” (*GAPDH* and *ACTIN*) and are shown as mean ± SEM. Data represent n = 3 independent biological experiments. Statistical significance was determined using one-way ANOVA assuming a lognormal distribution followed by Tukey’s multiple comparisons test. *P < 0.05, **P < 0.01, ***P < 0.001, **P < 0.0001; ns, not significant.

Analysis of circadian gene expression revealed distinct patterns between cbVG− and cbVG+ conditions. *ARNTL1* and *CLOCK* were modestly increased during infection, with no significant difference between cbVG− and cbVG+ conditions (**Figure 4C**). In contrast, *ARNTL2* and *NOCT* were strongly induced in cbVG+ infection (**Figure 4D–E**). All repressive and auxiliary circadian regulators (*PER1*, *NR1D1*, *NR1D2*, *BHLHE41*, and *KLF9*) were more strongly induced in cbVG+ than cbVG- infections (**Figure 4F**). Together, these data show that cbVG+ infection induces *ARNTL2, NOCT*, and multiple repressive clock components despite lower viral replication, whereas *ARNTL1* and *CLOCK* are not induced by cbVGs under these conditions.

To assess whether this cbVG-associated signature extends to a different negative-sense RNA virus, we examined expression of circadian genes in response to infection with respiratory syncytial virus (RSV), a member of a distinct family of mononegavirales which generates structurally diverse but functionally similar cbVG populations (1, 7, 65–67) (**Supplemental Figure 5A**). A549 cells were infected with cbVG-high (cbVG+) or cbVG-low (cbVG-) RSV stocks. Consistent with findings in SeV infection, *ARNTL2* was upregulated during infection, with cbVG+ cells showing a trend toward higher expression than cbVG− cells, although this difference did not reach statistical significance (P = 0.0762). *NOCT*, *NR1D1*, and *NR1D2* were more strongly induced in cbVG+ infection relative to both cbVG- and mock conditions (**Supplemental Figure 5B-C)** supporting a conserved cbVG-associated circadian transcriptional response across negative-sense RNA viruses.

### cbVG Reprograms circadian clock components by inducing two distinct viral sensing pathways

cbVGs activate RIG-I-like receptor/MAVS signaling and PKR signaling to induce type I and III interferon (IFN) production and protein translation shut off, respectively (5, 7, 9–14). To determine which mechanism drive the cbVG-specific circadian clock signature, and whether it depends on type I/III IFN signaling, we infected MAVS^KO^, PKR^KO^, and STAT1^KO^ A549 cells with cbVG− or cbVG+ SeV stocks and measured expression of circadian genes across the infection time course. Viral NP mRNA expression patterns between cbVG− and cbVG+ infections remained consistent across all cell lines, with cbVG− conditions consistently exhibiting higher replication (**Figure 5A** and **Supplemental Figure 6**). As expected, control A549 cells showed cbVG-specific induction of circadian clock targets, with *ARNTL2* and *NOCT* emerging as early responders (6 hpi, **Supplemental Figure 6).** In contrast, induction of the repressive clock program occurred later, with *NR1D1* becoming detectable by 12 hpi and *NR1D2*, and *PER1* showing cbVG-specific induction only at 24 hpi (**Figure 5 and Supplemental Figure 6**). These temporal differences suggest that distinct sensing events may drive early and late components of the circadian clock response.

**Figure 5.**
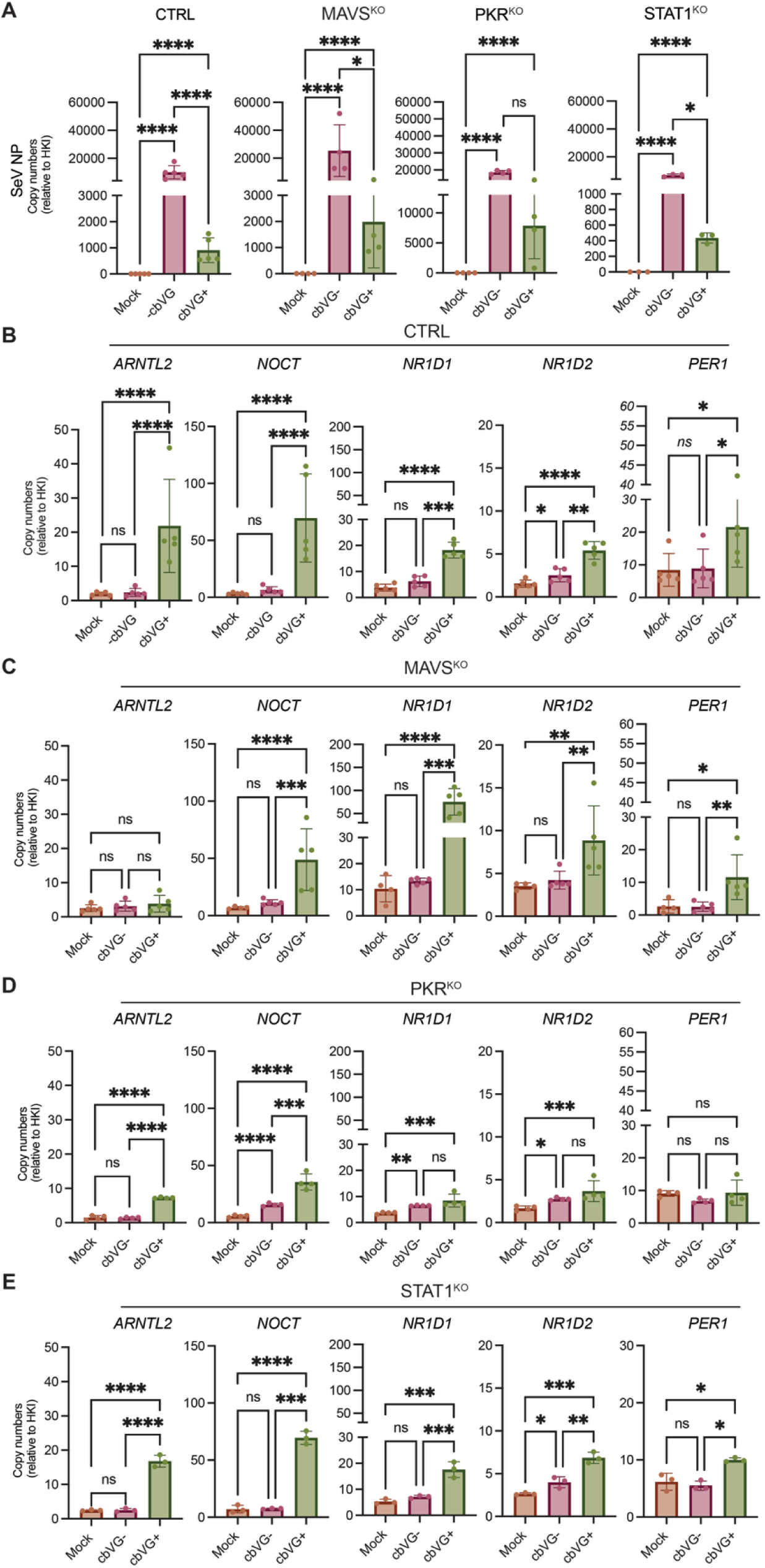
cbVG-driven induction of repressive circadian clock components and *ARNTL2* by distinct virus-sensing pathways. **(A)** *SeV NP* mRNA levels in CTRL, MAVS^KO^, PKR^KO^, and STAT1^KO^ A549 cells at 24 hpi under mock, cbVG−, and cbVG+ infection conditions, confirming viral replication across cell lines. **(B-E)** mRNA levels of circadian clock genes as measured by qPCR in **(B)** CTRL, **(C)** MAVS^KO^, **(D)** PKR^KO^ and **(E)** STAT1^KO^ A549 cells at 24 hpi under mock, cbVG−, and cbVG+ conditions. mRNA levels were normalized to a housekeeping gene index (HKI; *GAPDH* and *ACTIN*) and are shown as mean ± SEM. Data represent n = 5 (CTRL), n = 4 (MAVS^KO^), n = 4 (PKR^KO^), and n = 3 (STAT1^KO^) independent biological experiments. Statistical significance was determined using one-way ANOVA assuming a lognormal distribution followed by Tukey’s multiple comparisons test. *P < 0.05, **P < 0.01, ***P < 0.001, **P < 0.0001; ns, not significant.

We next looked at the impact of the sensing pathways on the expression of the broader panel of circadian genes at 24 hpi. Absence of MAVS selectively abolished cbVG-specific *ARNTL2* induction, whereas *NOCT*, *NR1D1*, *NR1D2*, and *PER1* remained inducible under cbVG-rich conditions (**Figure 5C**). Loss of PKR signaling produced a distinct pattern across the gene panel. *ARNTL2* cbVG-specific induction was decreased, but significant in PKR^KO^ cells, similar to *NOCT*. In contrast, *NR1D1*, *NR1D2* and *PER1* were no longer induced by cbVGs in PKR^KO^ cells, with both cbVG− and cbVG+ conditions becoming similarly elevated relative to mock at 24 hpi, indicating that PKR contributes to their induction in response to cbVG-enriched conditions rather than driving it directly (**Figure 5D**). MAVS-dependent induction of *ARNTL2* was further confirmed in an RSV infection, where infection-associated *ARNTL2* upregulation, including the trend toward higher expression in cbVG+ cells, was lost in MAVS-deficient cells (**Supplemental Figure 5D**).

Lastly, we looked at STAT1^KO^ cells to assess the contribution of STAT1 signaling to the induction of circadian clock genes. Despite the absence of STAT1, cbVG-associated induction of *ARNTL2*, *NOCT*, *NR1D1*, *NR1D2*, and *PER1* was maintained, indicating that activation of this circadian transcriptional program occurs independently of canonical IFN/STAT1 signaling (**Figure 5E**). Together, these findings identify two cellular sensing arms that differentially engage circadian clock components. MAVS signaling drives early *ARNTL2* induction, whereas PKR preferentially regulates the later induction of *PER1*, *NR1D1*, and *NR1D2*. *NOCT* regulation appears to involve additional pathways. All responses were independent of canonical IFN signaling.

### *Arntl2* overexpression broadly amplifies antiviral and inflammatory gene expression during SeV infection

Among the cbVG-induced circadian clock components identified, *ARNTL2* stood out as it is the only clock activating component selectively upregulated by cbVGs, mapped to a single virus sensing pathway, and with no previously characterized role during viral infection. Notably, its paralog *ARNTL1* which has known antiviral functions (35, 36, 38, 40, 68), was induced by infection independently of cbVG. To investigate the impact of *ARNTL2* on the host response to infection, we generated A549 cells stably overexpressing *ARNTL2* (BMAL2-A549). After confirming the overexpression by immunofluorescence (**Figure 6A**), the cells were infected with a cbVG− SeV stock. Because cbVG-rich infection induces endogenous *ARNTL2*, use of a cbVG-free stock allowed us to look at the consequences of *ARNTL2* overexpression during infection, independent on endogenous induction by cbVGs. Overexpression of *ARNTL2* in the absence of infection produced minimal transcriptional changes relative to control cells, confirming that the overexpression system itself does not broadly reprogram host gene expression at baseline (**Figure 6B; Supplemental Figure 7A–B**). Upon SeV infection, BMAL2-A549 cells showed a markedly broader transcriptional response relative to infected control cells, with predominantly upregulated differentially expressed genes (**Figure 6C**). Pathway enrichment identified interferon alpha response as the top upregulated pathway, followed by TNFα signaling via NF-κB, inflammatory response, complement, and apoptosis, while cell cycle pathways were suppressed (**Figure 6D**). qPCR validation confirmed significant upregulation of *OASL, TRIM22, CCL5, IFNB1, IFI27*, and *ISG15* in BMAL2-A549 cells relative to control cells infected under identical conditions (**Figure 6E**). Together, these findings show that elevated *ARNTL2* expression amplifies the transcriptional response to cbVG-free SeV infection, producing a broader antiviral and inflammatory response that is absent both in infected control A549 cells and in uninfected BMAL2-A549 cells. This transcriptional state therefore emerges during an infection with elevated *ARNTL2* levels, suggesting that cbVG-driven *ARNTL2* may repurposes its function to enhance the host antiviral response to the virus.

**Figure 6.**
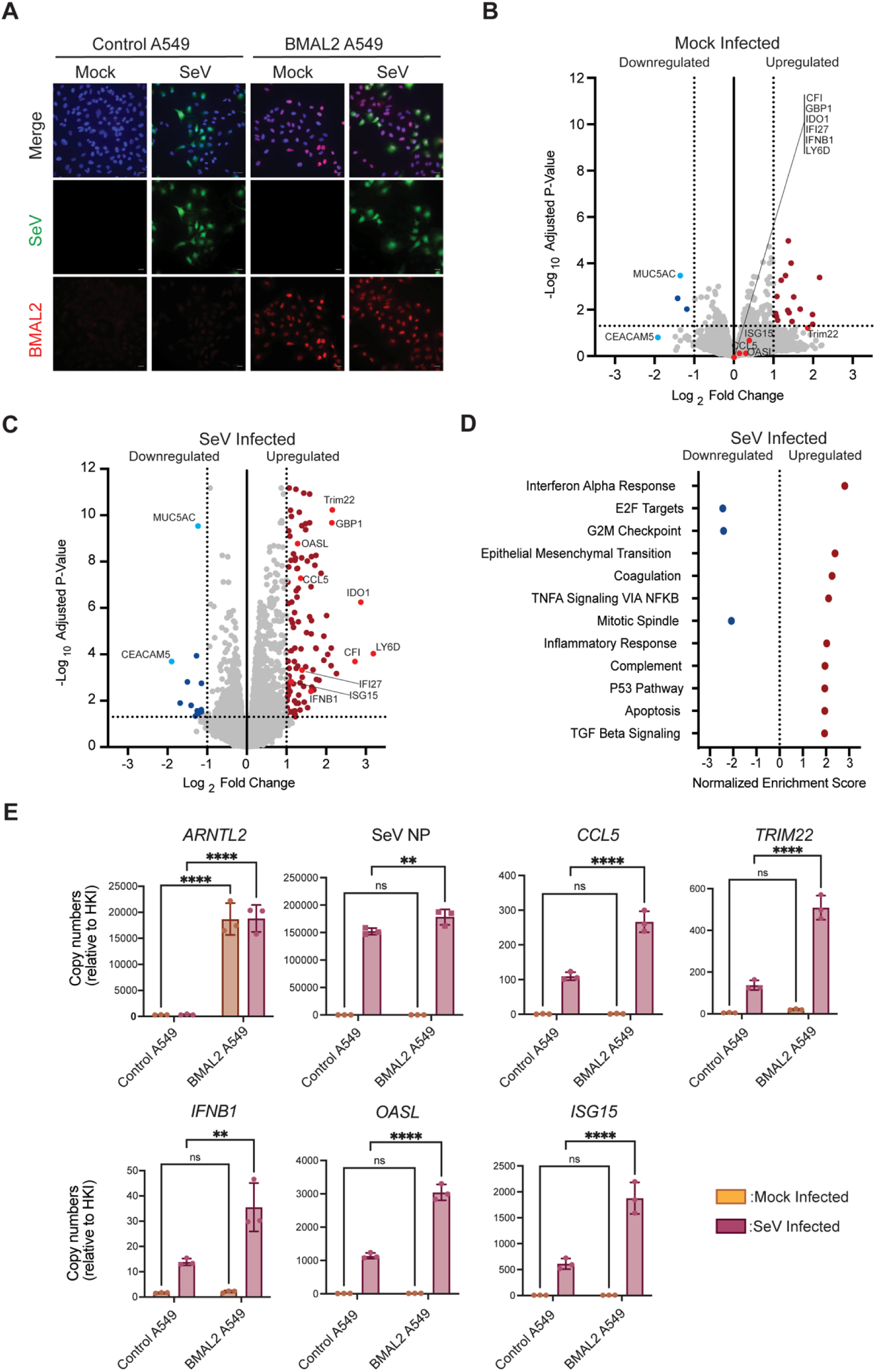
BMAL2 expression increases inflammatory gene expression during infection. A549 cells were transduced with a BMAL2-V5 tag expression plasmid. Control A549 and BMAL2 A549 cells were infected with SeV at an MOI of 1.5 TCID₅₀/cell using a cbVG-negative (cbVG−) stock. **(A)** At 24 hpi, cells were stained for V5 (red), SeV infection (eGFP, green), and nuclei (blue). **(B-D)** Cellular RNA was collected at 24 hpi and sequenced for host transcriptomic analysis. **(B)** Differential Gene Expression of mock-infected control A549 cells against BMAL2 A549 cells. **(C)** Differential Gene Expression of SeV-infected control A549 cells against BMAL2 A549 cells. **(D)** Functional Pathway Analysis of SeV-infected control A549 cells compared to BMAL2 A549 cells. **(E)** mRNA expression of SeV Nucleoprotein (NP), OASL, TRIM22, CCL5, IFNB1, IFI27, and ISG15 at 24 hpi as measured by qPCR. mRNA levels were normalized to housekeeping genes index “HKI” (*GAPDH* and *ACTIN*) and are shown as mean ± SEM. Data represent n = 3 independent biological experiments. Statistical significance was determined using one-way ANOVA assuming a lognormal distribution followed by Tukey’s multiple comparisons test. *P < 0.05, **P < 0.01, ***P < 0.001, **P < 0.0001; ns, not significant.

### *ARNTL2* knockdown selectively suppresses inflammatory genes during infection

To determine whether *ARNTL2* is functionally required for the observed amplified antiviral/inflammatory response (**Figure 6**), we depleted *ARNTL2* via siRNA knockdown in BMAL2-A549 cells and infected them with cbVG+ or cbVG- SeV (**Figure 7**). We used the BMAL2-A549 cells for these experiments as we could reliably track BMAL2 expression by immunostaining for the V5 tag. This approach allowed us to validate that a reduction in the V5 signal represents a reduction of BMAL2, as opposed to directly staining with the available antibodies against BMAL2 that in our hands either cross-react with other host proteins or don’t provide a clear signal for BMAL2. Immunofluorescence confirmed efficient *ARNTL2* knockdown (**Figure 7A**), with near-complete loss of the V5 signal in BMAL2 siRNA-treated cells relative to control siRNA. This was corroborated by reduced *ARNTL2* mRNA levels (**Figure 7C**). Upon infection, the viral nucleoprotein levels remained unchanged upon *ARNTL2* depletion (**Figure 7B**). *ARNTL2* knockout suppressed *CCL5* expression in both cbVG+ and cbVG-infections compared to control conditions (**Figure 7D**. TRIM22 expression showed similar trends, although significant suppression was only detected in cbVG- infection (**Figure 7E**). This differential sensitivity may reflect a more stable TRIM22 mRNA or a different threshold sensitivity for BMAL2 activity compounded with the fact that in cbVG+ infection there is residual viral-driven *ARNTL2* expression (**Figure 7C**). While the overexpression system limits direct assessment of physiological *ARNTL2* induction, these results establish *ARNTL2* as an enhancer of inflammatory response that is amplified during cbVG-rich infection. Altogether, our findings establish a direct link between viral genome composition and circadian clock reprogramming, reveal distinct MAVS- and PKR-dependent regulation of specific clock components, and identify *ARNTL2* as a possible modulator of innate inflammatory responses during infection.

**Figure 7.**
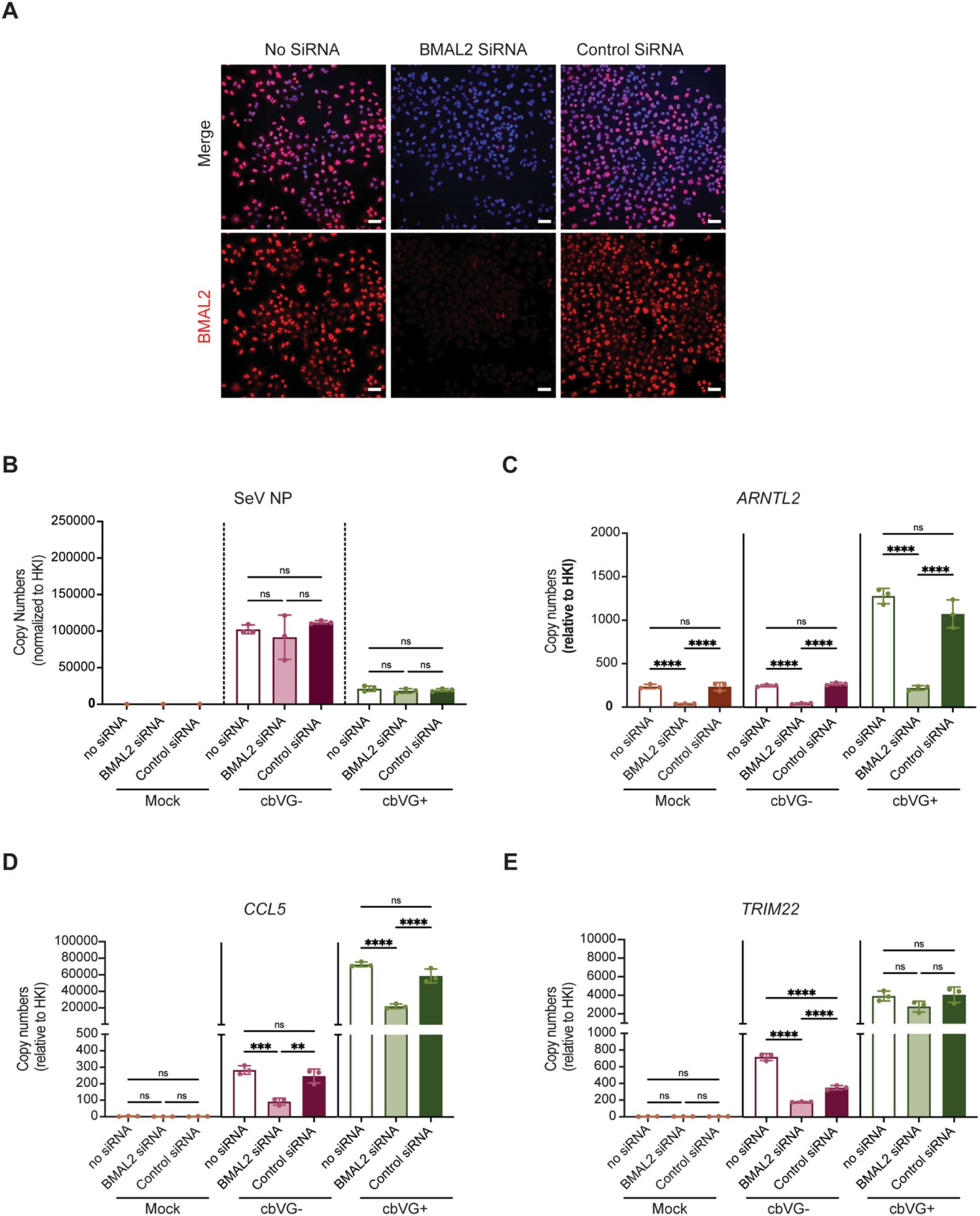
BMAL2 knockdown reduces inflammatory gene expression. BMAL2-expressing A549 cells were transfected with no siRNA, BMAL2 siRNA, or a control siRNA. **(A)** After 24 hours post transfection, cells were stained for BMAL2-V5 tag (red) and nuclei (blue). Scale bar :50 μm **(B-E)** At 24 hours post transfection, transfected A549 cells were infected with cbVG- SeV-eGFP or cbVG+ SeV-eGFP (20 HAUs) at an MOI of 1.5. Cellular RNA was collected at 24 hpi and gene expression was quantified for **(B)** SeV NP, **(C)** ARNTL2, **(D),** CCL5, and **(E)** TRIM22 by qPCR. mRNA levels were normalized to housekeeping genes index “HKI” (GAPDH and α-tubulin) and are shown as mean ± SEM. Data represent 3 biological replicates Statistical significance was determined using one-way ANOVA assuming a lognormal distribution followed by Tukey’s multiple comparisons test. *P < 0.05, **P < 0.01, ***P < 0.001, **P < 0.0001; ns, not significant.

## DISCUSSION

Circadian rhythms and viral infections are intimately connected, yet how the composition of the viral RNA population within an infected cell shapes the clock response has not been explored. Here we show that cbVGs, the immunostimulatory non-standard viral genomes generated during negative-sense RNA virus replication, induce a repression-biased clock transcriptional state in which MAVS- and PKR-dependent signaling regulate distinct but complementary components of the circadian response. Unexpectedly, this signature includes the selective induction of the circadian clock regulator *ARNTL2*. Our *ARNTL2* overexpression model (BMAL2-A549) demonstrated that elevated *ARNTL2* enhances antiviral and inflammatory transcriptional programs only during infection conditions, suggesting that its selective induction by cbVGs may actively reinforce, rather than simply accompany, the antiviral response.

We confirmed previous reports of A549 cells sustaining circadian-competency in culture (59) and further demonstrate that this property is maintained in infection-competent media. A549 cells are a lung epithelial cell line extensively used to model lung infections with common respiratory viruses, including influenza virus, parainfluenza viruses and RSV. By demonstrating A549’s circadian competency and identifying *ARNTL2* as a modulator of the host response to infection, our study establishes this cell line as a tractable model for studying virus-circadian interactions while minimizing the confounding effects of heterogeneous cell populations and multiple entrainment cues.

We also show that SeV infection reshapes rather than simply collapses the clock’s transcriptional organization, adding an important layer to the growing body of literature describing bidirectional interactions between viral infection and circadian biology. Previous studies have largely focused on how circadian state influences susceptibility to infection and antiviral immunity (34–38, 40, 41, 43, 47, 68–72), whereas considerably less is known about how infection feeds back onto the clock itself. Existing evidence indicates that microbial stimuli can reset circadian timing (73) and that viral factors can alter the expression of individual clock genes (44, 74, 75), supporting the idea that circadian programs remain responsive to pathogen-derived signals. Our findings extend these observations by demonstrating that infection-associated changes are not limited to individual clock components but involve a broader reorganization of the circadian clock transcriptional program. Importantly, this reorganization was most pronounced in cbVG-rich cells, where their presence was sufficient to drive the clock signature. This finding suggests that the circadian response to infection is influenced not only by infection itself, but also by the nature of the viral genomes present within infected cells. Given the established role of cbVGs in shaping antiviral immunity, these results reveal an additional layer through which viral genome heterogeneity can influence host-cell behavior.

Our data revealed that both the MAVS and the PKR virus-sensing pathways are involved in circadian clock disruption during infection. MAVS-dependent signaling is selectively required for *ARNTL2* induction, while PKR-dependent signaling drives *PER1* induction and shapes the cbVG specificity of *NR1D1* and *NR1D2* expression. Both arms operate independently of STAT1, demonstrating that the cbVG-specific clock signature reflects direct innate sensing rather than a secondary consequence of interferon-driven ISG amplification. As cbVGs are robust triggers of the MAVS and PKR pathways leading to the induction of the host antiviral response during SeV and RSV infection, their impact on the circadian clock suggests that clock components are interconnected with the cell ability to response to infection. Consistent with this possibility, infected cells progressively accumulated within restricted regions of circadian pseudotime. Notably, the population in which cbVGs first emerged remained enriched within the same early circadian phase interval across successive infection time points, raising the untested possibility that infection alters the normal progression through circadian state to favor host defense.

Among the genes induced by cbVGs, *ARNTL2* occupies a unique position. It is the only activating circadian clock component whose expression is selectively increased during cbVG-rich infection, its induction is conserved across two viruses, and it maps specifically to MAVS-dependent signaling. The function of its encoded protein BMAL2 during viral infection was entirely uncharacterized. *ARNTL2* is a paralog of *ARNTL1* that forms heterodimers with *CLOCK* and *NPAS2* to drive E-box transcription; whether it can functionally substitute for *ARNTL1* under conditions of *ARNTL1* loss remains debated (76–79). During non-viral inflammatory responses, *ARNTL2* expression was reported to be induced by TNFa via NF-κB signaling in primary human fibroblasts, where the *ARNTL2/NPAS2* heterodimer drives E-box transcription with distinct target gene selectivity compared to the canonical *ARNTL1/CLOCK* complex (80). Our findings suggest that this alternative circadian regulatory program may also be engaged during RNA virus infection. Together with the additional clock regulators identified in our study, these observations raise the possibility that antiviral innate immune signaling engages a distinct circadian transcriptional network rather than simply perturbing the canonical circadian clock. Defining the components and function of this infection-responsive circadian program may provide new insight into how circadian regulation is integrated with antiviral immunity.

Interestingly, *ARNTL2* has been proposed to compensate for *ARNTL1* under conditions of *ARNTL1* loss (76), our data argue against this interpretation. At 24 hpi, *ARNTL1* transcript levels were induced to comparable levels in both cbVG− and cbVG+ conditions, indicating that *ARNTL2* induction reflects a cbVG-induced transcriptional program rather than a response to the activator depletion. Whether this distinction holds at the protein level remains an open question. PKR-dependent translational shutoff in cbVG-high cells may differentially constrain *ARNTL1* and *ARNTL2* protein accumulation despite equivalent mRNA, a possibility that will require protein-level characterization to resolve.

Overexpression of *ARNTL2* had minimal transcriptional effect at baseline. Upon infection, however, *ARNTL2*-overexpressing cells showed a markedly broader antiviral and inflammatory transcriptional response, including interferon response, TNFα/NF-κB signaling, inflammatory response, and complement programs, that were absent in infected control cells. This transcriptional state emerges specifically at the intersection of *ARNTL2* upregulation and active infection, indicating that its activity is context dependent. Knockdown of endogenous *ARNTL2* in infected cells suppressed *CCL5* and *TRIM22* expression, confirming functional role for BMAL2. The mechanism by which BMAL2 facilitates the response to infection remains unknown. Notably, *ARNTL1* silencing has been shown to enhance antiviral gene expression in lung epithelial cells and restrict SARS-CoV-2 (38). However, ARNTL1 expression was not differentially regulated by cbVG accumulation in our system, whereas *ARNTL2* was selectively induced in cbVG-rich cells. These observations raise the possibility that *ARNTL2* participates in some antiviral regulation through mechanisms that are distinct from, or only partially overlapping with, those of *ARNTL1*, a question that will require further mechanistic investigation.

The functional consequences of a repression-biased clock state in cbVG-high cells also remain to be determined. In other inflammatory contexts, *NR1D1* suppresses cytokine production and dampens immune activation (81–83), but whether its induction in our experimental conditions serves a similar function, represents a compensatory response within a dysregulated clock, or reflects an independent consequence of PKR signaling is unknown. Furthermore, the temporal dynamics of circadian clock reprogramming and its relationship to cbVG accumulation remain unexplored. Whether pre-existing circadian state influences the generation of cbVGs, or whether cbVG accumulation progressively reshapes circadian organization to reinforce antiviral responses, remains unresolved. Dissecting this bidirectional relationship will be essential to understanding how circadian regulation and viral genome heterogeneity influence one another throughout infection.

Taken together, this work establishes viral genome heterogeneity as a previously unrecognized axis of host circadian regulation during infection and provides evidence of an intricate circuit between circadian clock genes and the response to an infection. These observations open new avenues on how we think about the host response to negative-sense RNA viruses.

## MATERIALS AND METHODS

### Cell lines

A549 human lung epithelial cells (ATCC CCL-185), including wild-type (WT), CRISPR control (CTRL), MAVS knockout (MAVS^KO^), PKR knockout (PKR^KO^), and STAT1 knockout (STAT1^KO^) derivatives, were maintained in Dulbecco’s Modified Eagle Medium (DMEM; Thermo Fisher) supplemented with 10% fetal bovine serum (FBS) (Milipore Sigma), 2 mM L-glutamine (Invitrogen), 1 mM sodium pyruvate (Invitrogen), and 50 μg/mL gentamicin (Thermo Fisher) at 37°C in a humidified incubator containing 5% CO2. The CRISPR-engineered cell lines were kindly provided by Dr. Susan Weiss (University of Pennsylvania) and have been previously described and validated (7, 8, 84). An *ARNTL2*-overexpressing A549 cell line (A549-BMAL2) was generated by transduction with the human *ARNTL2* expression construct TFORF3265 (Addgene plasmid #144741deposited by Feng Zhang) (85). Stable cell lines were established by antibiotic selection and validated by assessment of *ARNTL2* expression. Cell cultures were routinely tested for mycoplasma contamination using a PCR-based assay according to our standardized laboratory protocol (https://dx.doi.org/10.17504/protocols.io.81wgbxw71lpk/v1).

### Viruses

Sendai virus (SeV) strain Cantell enriched in copy-back viral genomes (cbVG-high), used in the previously published datasets reanalyzed here, was generated in 10-day-old embryonated chicken eggs as described (66). For details see our published protocol: https://www.protocols.io/edit/growing-sev-in-eggs-deam3ac6). Experimental infections performed for this study used recombinant SeV Cantell viruses engineered to express fluorescent reporters. Circadian gene expression and pathway analyses (Figures 4–5 and Supplemental Figures 5–6) used rSeVC-LmiRFP670, which expresses miRFP670 fused to the C-terminus of the viral L protein and was generated using the SeV Cantell reverse genetics system as previously described (86). Briefly, the full-length plasmid pSL1180-rSeVC-LmiRFP670 was constructed by inserting the miRFP670 coding sequence downstream of the L gene through a GRP linker while maintaining compliance with the rule of six. Recombinant virus was rescued by co-transfection of the full-length viral genome plasmid with helper plasmids encoding NP, P, and L proteins into BSR-T7/5 cells using Lipofectamine LTX with Plus Reagent (Invitrogen). Virus-containing lysates were amplified in 10-day-old embryonated chicken eggs for 40 h at 37°C, and viral stocks were harvested from allantoic fluid and titrated on LLC-MK2 cells. *ARNTL2* overexpression and siRNA knockdown experiments (**Figures 6-7**) used a recombinant SeV-eGFP reporter virus, also derived from the SeV Cantell strain, to enable visualization of infected cells by fluorescence microscopy. Virus rescue, propagation, amplification, and titration were performed as previously described (86) and according to our detailed published protocol (https://dx.doi.org/10.17504/protocols.io.dm6gpzeojlzp/v1). Respiratory syncytial virus (RSV) stocks were propagated in HEp-2 cells as previously described and according to our published protocol (https://www.protocols.io/view/rsv-expansion-and-titration-5qpvok39xl4o/v1). RSV stocks enriched or depleted in copy-back viral genomes (cbVG-high and cbVG-low, respectively) were generated and characterized as previously reported (87); briefly, viral populations were serially passaged under conditions promoting cbVG accumulation or depletion, and cbVG content was confirmed by sequencing and quantitative analysis (87).

### Viral Infections

Virus infections were performed according to standardized laboratory protocols (Protocols.io: https://dx.doi.org/10.17504/protocols.io.dm6gpzeojlzp/v1). For SeV experiments, cells were infected with recombinant SeV Cantell reporter viruses at a multiplicity of infection (MOI) of 1.5 TCID_₅₀_/cell. cbVG− conditions consisted of reporter virus alone, whereas cbVG+ conditions were generated by supplementing the same viral inoculum with 20 hemagglutination units (HAU) of purified SeV cbVG particles, as previously described (88). Purified cbVG particles were isolated from the allantoic fluid of SeV-infected embryonated chicken eggs by density ultracentrifugation through a 5–45% sucrose gradient (88). For RSV experiments, cells were infected with cbVG-low or cbVG-high RSV stocks at an MOI of 3 for 2 h at 37°C, followed by replacement with fresh medium containing 2% FBS.

### Single-cell RNA-sequencing data analysis

Single-cell RNA-sequencing data were obtained from Yang et al. (54) which profiled human A549 cells under mock and SeV infection conditions at 6, 12, and 24 hours post-infection. The libraries captured dual transcriptome, enabling the simultaneous quantification of host genes and viral RNA. Quality control and initial preprocessing were performed as described (54). Specifically, host transcriptome was processed using 10X Genomics CellRanger 7.2.0 and R Seurat package (89), and cells producing copy-back viral genomes (cbVGs) were identified using the VODKA2 pipeline (60), with these metrics integrated into the single-cell metadata. To ensure comparability across timepoints and conditions while balancing statistical power, 5,000 high-quality cells were randomly selected from each sample. Manifold visualization was performed using Uniform Manifold Approximation and Projection (UMAP). Transcriptional clusters were identified using a shared nearest neighbor (SNN) modularity optimization-based clustering algorithm at a resolution of 0.1, specifically to isolate interferon (IFN)-producing populations from the general infected population. Differential gene expression analysis between clusters was performed using the Wilcoxon rank-sum test, requiring genes to be expressed in at least 25% of cells and exhibit a minimum log2 fold-change of 0.25. Cell cycle phase assignment was performed using Seurat’s CellCycleScoring function (89) with the updated 2019 human cell cycle gene sets (cc.genes.updated.2019), which assign cells to G1, S, or G2M phases based on expression of phase-specific marker genes (90). All analyses were performed in R version 4.4.2. Visualizations were generated using ggplot2 (91).

### Circadian phase inference

We adapted the principal component-based phase inference method developed for cell cycle analysis by Schwabe et al. (52) to estimate circadian phase from clock gene expression. Phase inference was restricted to a set of 23 core circadian clock genes (*ARNTL1, ARNTL2, CLOCK, PER1–3, CRY1–2, NR1D1–2, RORA–C, DBP, TEF, NPAS2, BHLHE40–41, CIART, NOCT, KLF9, CSNK1D–E*), extracted from the normalized expression matrix. We limited the input to the core transcriptional-translational feedback machinery and its interlocking loops (15), rather than the broader set of rhythmically expressed clock-controlled output genes, as these core components have conserved, well-defined phase relationships that provide a stable reference for phase inference. Gene expression values were centered and scaled to unit variance prior to PCA. Circadian phase (θ) for each cell was calculated as θ = atan2(PC2, PC1) and converted to a 0–2π range. Cells were binned into 28 equal-width phase bins, and mean expression was calculated per gene per bin. Expression trajectories were visualized by LOESS smoothing (span = 0.7) of the binned data.

### Circadian luciferase lentivirus production

Lentiviral reporters expressing firefly luciferase driven by *ARNTL1* (previously described in Liu et al., 2008; Zhang et al., 2009) or *PERIOD2* (previously described in Ramanathan et al., 2012) promoters were generated in the laboratory using plasmids (gift from Dr. Andrew Liu, University of Memphis) and packaging plasmids psPAX2 and pMD2.G (gift from Didier Trono (Addgene plasmid # 12260 and 12259). Hek293T/17 cells (ATCC) were plated at 3 million cells per 100 mm dish in complete DMEM media with 10% FBS (Thermo Fisher) and 5% Pen/Strep (Thermo Fisher) in a 5% CO_2_ incubator for 24-30 hours. Before transfection, media was changed to DMEM with 5% FBS (Fisher) without antibiotics. For transfection, cells were incubated with Fugene 6 (Promega) and plasmids for 12 hours, then changed to complete media. Viral particles in media were collected after 48 and 72 hours, centrifuged (RT, 3 min at 0.4 rcf) and filtered through a PES 0.45 μm filter (Fisher). The presence of viral particles was confirmed with Lenti-X GoStix Plus test (Takara).

### Lentiviral transductions with the luciferase reporters for *ARNTL1* and *PER2*

A549 cells were transduced with lentiviral reporters expressing firefly luciferase driven by ARNTL1 or *PERIOD2*. Cells were grown in T-25cm^2^ flasks for 24 hours and incubated for 10 minutes at 37°C in 3mL complete media supplemented with 15μg polybrene (Millipore). Following incubation, 500μL of virus stock was added to each culture. Media was changed after 24 h at 37°C. Infected cells were selected using blasticidin (1.25 μg/mL, Thermo Fisher), and luciferase expression was confirmed by recording bioluminescence *in vitro*.

### Bioluminescence recordings and analysis

A549 cells were plated in 35 mm (BD Falcon, Fisher) petri dishes, supplemented with cell culture media containing 0.1mM D-luciferin (Goldbio), sealed with vacuum grease, and placed in a light-tight 36°C incubator containing photo-multiplier tubes (PMTs) (Hamamatsu Photonics). Each dish was placed under one PMT and bioluminescence was recorded as photons per 180 seconds for 96 hours. Bioluminescence data was detrended with a 24-hour moving average and analyzed in ChronoStar 1.0. The correlation coefficient (CC) of a best-fit circadian cosine function was calculated using ChronoStar 1.0 to assess circadian rhythmicity in A549 cells. CC values above 0.7 were considered circadian.

### RNA extraction, cDNA synthesis, and quantitative PCR

Total RNA was extracted using the MagMAX mirVana Total RNA Isolation Kit (Thermo Fisher Scientific, #A27828) on a KingFisher Flex automated purification system according to the manufacturer’s instructions and our standardized published protocol (https://www.protocols.io/edit/for-tissue-cells-and-tissue-samples-rna-extraction-detx3epn). cDNA was synthesized from 150 ng of total RNA using the High-Capacity RNA-to-cDNA Kit (Thermo Fisher) following our published protocol (https://www.protocols.io/edit/reverse-transcription-for-qpcr-ssiii-deqe3dte). Quantitative PCR (qPCR) was performed using Power SYBR Green PCR Master Mix (Thermo Fisher) and gene-specific primers (Supplementary Table S5) according to our published protocol (https://dx.doi.org/10.17504/protocols.io.8epv5rxw6g1b/v1). Relative gene expression was calculated using the ΔΔCt method and normalized to the geometric mean of *GAPDH* and *ACTB* expression(92). All reactions were performed in technical triplicate, and technical replicates were averaged prior to statistical analysis of biological replicates.

### Bulk RNA sequencing *ARNTL2*-overexpressing A549

Control and *ARNTL2*-overexpressing A549 cells were either mock-treated or infected with SeV at an MOI of 1.5. Cellular RNA harvested at 24 hpi and submitted to Plasmidsaurus (Eugene, OR, USA) for bulk RNA sequencing. Libraries were prepared using the Plasmidsaurus RNA-Seq workflow, which employs a 3′ mRNA counting strategy for transcriptome-wide gene expression profiling, and sequenced on an Illumina platform to a depth of approximately 10 million reads per sample. Raw sequencing data were processed to generate gene-level count matrices, which were used for downstream differential gene expression and pathway enrichment analyses.

### *ARNTL2* siRNA-mediated knockdown

*ARNTL2*-overexpressing A549 cells were transfected with ON-TARGETplus SMARTpool siRNA targeting human *ARNTL2* (Horizon Discovery/Revvity; Entrez Gene ID: 56938) or ON-TARGETplus Non-targeting Control siRNA according to the manufacturer’s instructions and our published protocol (https://www.protocols.io/edit/transfection-protocol-dem43c8w). ON-TARGETplus SMARTpool reagents consist of four target-specific siRNAs combined into a single pool to maximize knockdown efficiency while minimizing off-target effects. Cells were transfected using DharmaFECT 3 (Dharmacon, T-2003) at a final siRNA concentration of 20 nM. Knockdown efficiency was confirmed by RT-qPCR and immunofluorescence 24 h post-transfection before infection.

### Immunofluorescence

Immunofluorescence staining was performed according to our standardized published protocol (https://www.protocols.io/view/immunofluorecence-assay-ifa-rm7vzjxz4lx1/v1). Briefly, cells were fixed with 4% paraformaldehyde, permeabilized with 0.2% Triton X-100, and incubated sequentially with primary and fluorophore-conjugated secondary antibodies. Nuclei were counterstained with Hoechst 33342, and coverslips were mounted using ProLong Diamond Antifade Mountant (Thermo Fisher). Primary antibodies used in this study included anti-SeV nucleoprotein (NP; clone M73/2, a gift from Alan Portner), anti-BMAL2 (Sigma-Aldrich), and anti-V5 (Invitrogen).

## Supporting information

All supplementary figures

## DATA AVAILABILITY

Raw sequencing data from *ARNTL2*-overexpressing A549 cells generated in this study were deposited in the NCBI Gene Expression Omnibus (GEO; accession GSE337458). Publicly available sequencing data from the study by Yang et al. were downloaded from the NCBI Sequence Read Archive (SRA; BioProject accession PRJNA1390230), and gene expression data from the study by Xu et al. were obtained from the NCBI Gene Expression Omnibus (GEO; accession GSE96774). All other data will be available at ImmPort.

## ACKNOWLEDGMENTS

We thank Dr. Susan Weiss for providing the A549 PKR knockout, MAVS knockout, STAT1 knockout, and parental control A549 cell lines. We also thank Ana Ruth Tronco for her assistance with experimental procedures and sample processing for the experiments presented in Figures 4 and 5. This study was supported by the BJC Investigator Program (CBL) and NIH grants AI137062 (to CBL), NS134885 (to EH), and T32-AI007172 (to NRE).

## AUTHOR CONTRIBUTIONS

Conceptualization: N.R.E., E.A., C.B.L.

Methodology: N.R.E., E.A., M.H., M.F.G-A., E.D.H., C.B.L.

Validation: N.R.E., E.A., M.H., M.F.G-A.

Formal analysis: N.R.E., E.A., M.H.

Investigation: N.R.E., M.H., M.F.G-A.

Resources: Y.Y., E.D.H., C.B.L.

Data curation: N.R.E., E.A., M.H. C.B L

Writing-original draft: N.R.E., E.A., C.B.L.

Writing-review & editing: N.R.E., E.A., M.H., M.F.G-A., Y.Y., E.D.H., C.B.L.

Supervision: C.B.L. Funding acquisition: C.B.L.

