## Supplementary material for "Circadian Clock Gene Modulation and Selective Reprogramming in Response to Virus Infection": All supplementary figures

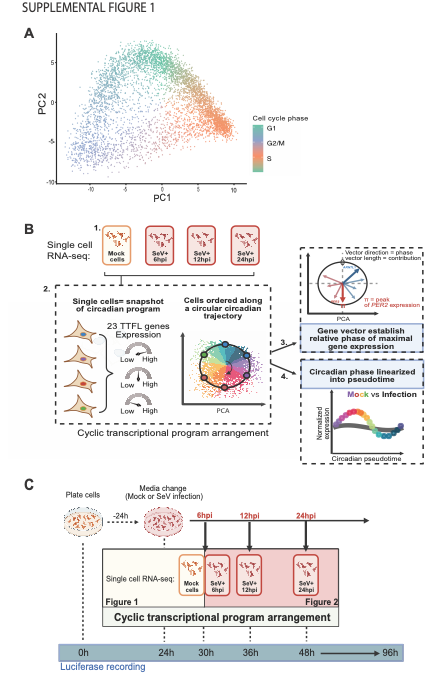

**Supplemental Figure 1. Validation of PCA-based trajectory reconstruction and single-cell experimental workflow. A)** Principal component analysis (PCA) trajectory reconstruction of cell-cycle progression in uninfected mock A549 cells. Individual cells are projected onto a PC1–PC2 coordinate space using a curated cell-cycle gene panel, with each cell colored by its canonical cell-cycle phase: G1(green), G2/M(blue), or S phase (orange). Unsynchronized cells distribute into a continuous, arc-like cyclical trajectory matching sequential biological progression through cell division, validating that the PCA framework accurately resolves cyclic host transcriptional programs. **(B)** Schematic workflow for single-cell transcriptomic profiling and circadian phase inference. **1.** Mock-treated and Sendai virus (SeV)-infected (MOI 1.5) A549 cells were collected at 6, 12, and 24 hours post-infection (hpi) for single-cell RNA sequencing (scRNA-seq). **2.** Because individual cells capture different points within the circadian transcriptional program, the expression profiles of 23 TTFL genes were used to reconstruct a continuous circadian trajectory by principal component analysis (PCA). **3.** Gene loading vectors were used to infer the relative phase of maximal gene expression and orient the reconstructed trajectory. **4.** The circular trajectory was subsequently linearized into circadian pseudotime (θ), enabling quantitative comparison of circadian gene expression dynamics between mock and SeV-infected cells. **(C)** Experimental timeline for the single-cell transcriptomic sampling and the continuous circadian bioluminescence recordings. A549 cells were plated for 24 hours before media change, at which point cells were either mock-treated or infected with SeV and sampled for single-cell RNA sequencing at 6, 12, and 24 hours post-infection (30, 36, and 48 hours after plating, respectively). These collection timepoints correspond to defined positions within the continuous *PER2*::LUC and *ARNTL1*::LUC bioluminescence recordings shown in Figure 1, providing an independent circadian reference for interpreting the reconstructed single-cell circadian trajectory throughout infection.

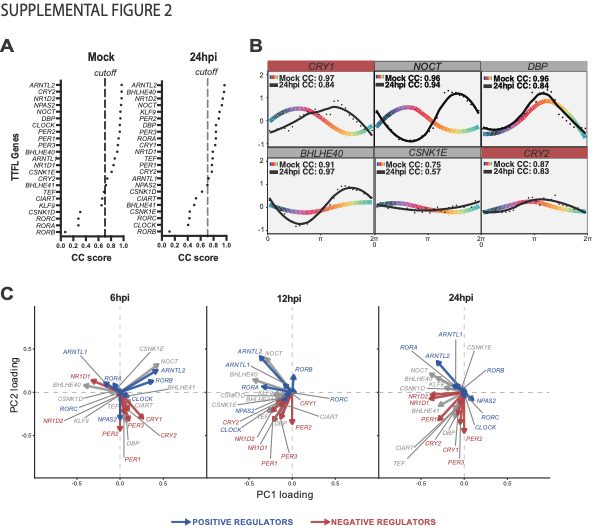

**Supplemental Figure 2.** **Sendai virus infection progressively reorganizes host circadian clock transcriptional structure and reduces activator coherence.**  **(A)** Cosine fit correlation coefficient (CC) score distributions for 23 core clock genes in Mock (left) and 24hpi (right) cells, ordered hierarchically by fit accuracy. Vertical dashed lines indicate stringency cutoffs used to identify high-coherence, rhythmically cycling transcripts (CC ≥ 0.70) **(B)** Single-cell expression profiles and corresponding sinusoidal model fits for clock components selected due to their high baseline CC scores in the mock condition (*CRY1, NOCT, DBP, BHLHE40, CSNK1E, CRY2*), plotted against inferred circadian phase (0 to 2π radians). Dots and colored curves track uninfected cells (Mock); solid black curves represent fitted trajectories at 24hpi. Calculated CC scores are annotated within each panel. **(C)** PCA loading plots showing dynamic shifts in gene-specific weights across the rotated PC1–PC2 coordinate space at 6, 12, and 24 hpi. Vectors indicate the direction and magnitude of contribution for core positive regulators (blue), core negative regulators (red), and auxiliary/output transcripts (gray). Vector realignment reveals a coordinated structural reorganization of the clock machinery over time.

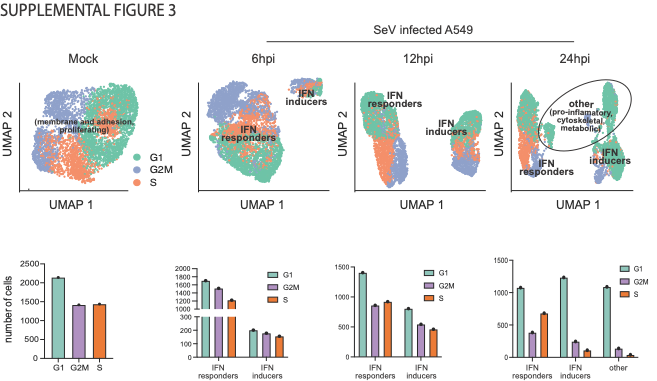

**Supplemental Figure 3.** **Transcriptomic profiling of Sendai virus-induced cell states reveals host defense clusters that emerge independently of cell-cycle phase.** Uniform Manifold Approximation and Projection (UMAP) plots (top) showing cell-cycle phase assignments (G1, green; S, orange; G2/M, blue) in mock cells and at 6, 12, and 24 hours post-infection (hpi) with SeV. Corresponding bar plots (bottom) show the distribution of cells across cell-cycle phases within each identified cluster over the course of infection, illustrating progressive changes in cell-cycle composition associated with the emergence of distinct infection-induced cellular states.

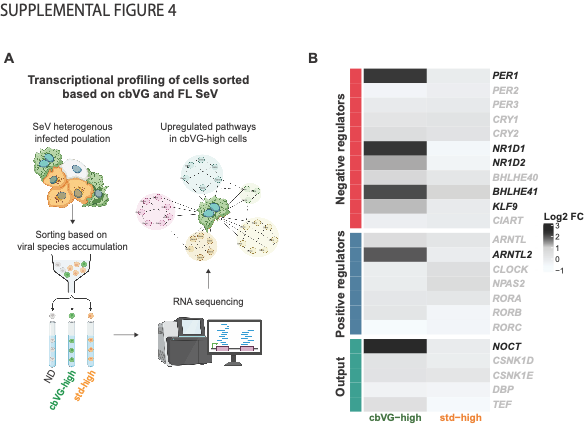

**Supplemental Figure 4. Sorting by viral genome composition reveals differential circadian gene expression in cbVG-high cells. (A)** Workflow for sorting SeV-infected A549 cells based on viral RNA accumulation into cbVG-high, standard genome-high (std-high), and non-detected (ND) populations, followed by RNA sequencing at 24 hpi. **(B)** Heatmap showing log2 fold change (log2FC) of circadian clock genes in cbVG-high and FL-high cells relative to ND. Genes are grouped by functional class: negative regulators, positive regulators, and output/regulatory modulators. Bold gene names indicate genes with log2FC ≥ 1 in cbVG-high cells. Data represent n = 2 independent biological replicates per condition.

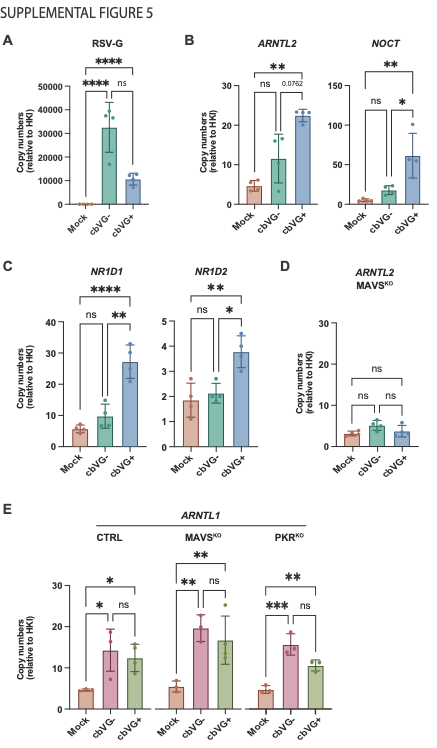

**Supplemental Figure 5. RSV infection recapitulates the cbVG-associated circadian transcriptional signature and supports MAVS-dependent regulation of ARNTL2.** **(A)** RSV-G mRNA levels in A549 cells at 24 hpi under mock (Mock), low-cbVG-dose (cbVG-), and high-cbVG-dose (cbVG+) RSV infection conditions. **(B–C)** mRNA levels of circadian clock genes ARNTL2 and NOCT (activating component and clock-controlled output, respectively) and repressive regulators PER1, NR1D1, and NR1D2 in A549 cells at 24 hpi. **(D)** ARNTL2 mRNA levels in MAVS^KO^ A549 cells at 24 hpi under mock, cbVG-, and cbVG+ RSV infection conditions. **(E)** ARNTL1 mRNA levels in CTRL, MAVS^KO^, and PKR^KO^ A549 cells at 24 hpi under mock, cbVG−, and cbVG+ SeV infection conditions. mRNA levels were normalized to a housekeeping gene index (HKI; *GAPDH* and *ACTIN*) and are shown as mean ± SEM. Data represent n = 4 (panel A-D) and n = 3-4 (panel E) independent biological experiments. Statistical analysis was performed using one-way ANOVA with multiple comparisons (*p < 0.05, **p < 0.01, ****p < 0.001; ns, not significant).

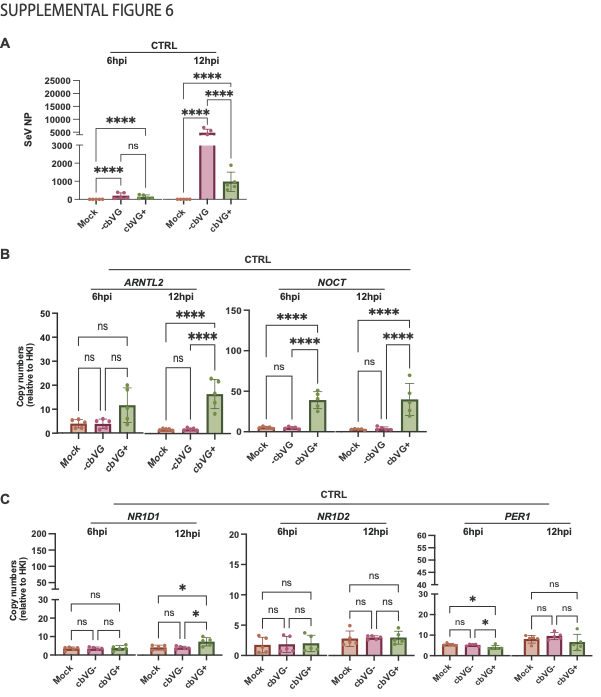

**Supplemental Figure 6. cbVG-driven clock gene induction follows distinct early and late kinetics in CTRL A549 cells (A)** SeV NP mRNA levels in CTRL A549 cells at 6 and 12 hpi under mock (M), cbVG− and cbVG+ infection conditions, confirming viral replication across cell lines and timepoints. **(B–D)** mRNA levels of circadian clock genes ARNTL2, NOCT, (B), NR1D1, NR1D2, and PER1 in CTRL A549 cells at 6 and 12 hpi under mock, cbVG−, and cbVG+ conditions. mRNA levels were normalized to a housekeeping gene index (HKI; *GAPDH* and *ACTIN*) and are shown as mean ± SEM. Data represent n = 5 independent biological experiments. Statistical significance was determined using one-way ANOVA assuming a lognormal distribution followed by Tukey's multiple comparisons test. *P < 0.05, **P < 0.01, ***P < 0.001, ****P < 0.0001; ns, not significant.

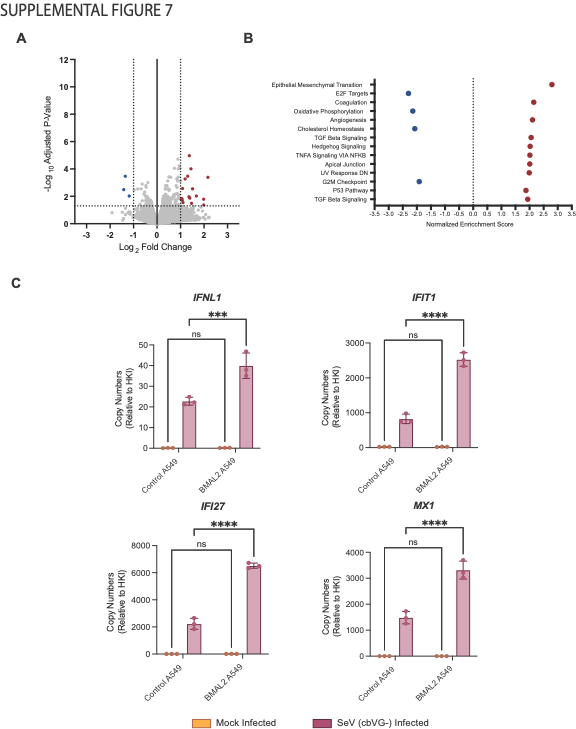

**Supplemental Figure 7. ARNTL2-driven enhancement of innate immune gene expression requires active SeV infection (A)** Volcano plot showing differential gene expression between mock-infected control A549 and BMAL2-overexpressing A549 cells. Significantly upregulated genes (log₂ fold change > 1, adjusted p < 0.05) are shown in red; significantly downregulated genes (log₂ fold change > 1, adjusted p < 0.05) in blue. **(B)** Gene Set Enrichment Analysis of mock-infected control A549 compared to BMAL2-overexpressing A549 cells, showing normalized enrichment scores for the top enriched (red) and depleted (blue) Hallmark pathways. **(C)** mRNA expression of innate immune genes IFNL1, IFIT1, IFI27, and MX1 in control A549 and BMAL2-overexpressing A549 cells under mock and SeV-infected conditions at 24 hpi. mRNA levels were normalized to a housekeeping gene index (HKI; *GAPDH* and *ACTIN*) and are shown as mean ± SEM. Data represent n = 3 independent biological experiments. Statistical significance was determined using one-way ANOVA assuming a lognormal distribution followed by Tukey's multiple comparisons test. *P < 0.05, **P < 0.01, ***P < 0.001, ****P < 0.0001; ns, not significant.

**Table S1.** Top 10 differentially expressed genes per cluster in mock sample.

| p_val | avg_log2FC | pct.1 | pct.2 | p_val_adj | cluster | gene | annotation |
| --- | --- | --- | --- | --- | --- | --- | --- |
| 7.68E-44 | 1.811 | 0.383 | 0.210 | 2.03E-39 | 0 | AL356124.1 | Membrane and adhesion (Homeostatic / Structural) |
| 2.61E-79 | 1.552 | 0.506 | 0.258 | 6.90E-75 | 0 | TMEM150A |  |
| 1.59E-40 | 1.517 | 0.379 | 0.209 | 4.20E-36 | 0 | RBMS3 |  |
| 7.90E-31 | 1.510 | 0.261 | 0.123 | 2.09E-26 | 0 | ST6GAL1 |  |
| 3.98E-47 | 1.501 | 0.485 | 0.306 | 1.05E-42 | 0 | PLCXD3 |  |
| 1.78E-30 | 1.464 | 0.332 | 0.189 | 4.72E-26 | 0 | TRPC6 |  |
| 1.81E-31 | 1.451 | 0.321 | 0.182 | 4.78E-27 | 0 | ITGB8 |  |
| 1.18E-32 | 1.381 | 0.504 | 0.364 | 3.12E-28 | 0 | FGFBP1 |  |
| 1.45E-37 | 1.340 | 0.411 | 0.252 | 3.82E-33 | 0 | SETBP1 |  |
| 1.70E-22 | 1.296 | 0.493 | 0.394 | 4.50E-18 | 0 | GACAT2 |  |
| 0 | 6.923 | 0.439 | 0.008 | 0 | 1 | PIF1 | Proliferating (S / G2/M phase) |
| 7.44E-290 | 4.499 | 0.531 | 0.089 | 1.97E-285 | 1 | HIST1H1B |  |
| 5.37E-243 | 3.854 | 0.778 | 0.469 | 1.42E-238 | 1 | HIST1H1D |  |
| 0 | 3.802 | 0.999 | 0.566 | 0 | 1 | UBE2C |  |
| 2.21E-292 | 3.789 | 0.706 | 0.231 | 5.84E-288 | 1 | HIST1H2AI |  |
| 2.22E-305 | 3.728 | 0.669 | 0.191 | 5.88E-301 | 1 | HIST1H2AH |  |
| 0 | 3.689 | 0.990 | 0.283 | 0 | 1 | AURKB |  |
| 1.10E-141 | 3.667 | 0.254 | 0.025 | 2.92E-137 | 1 | HIST1H2AL |  |
| 3.46E-291 | 3.510 | 0.622 | 0.140 | 9.15E-287 | 1 | HIST2H2BF |  |
| 0 | 3.500 | 0.894 | 0.174 | 0 | 1 | KIF2C |  |

p_val: unadjusted p-value; avg_log2FC: average log2 fold change in expression between the indicated cluster and all remaining cells; pct.1 and pct.2: percentage of cells expressing the gene within the indicated cluster and all other cells, respectively; p_val_adj: Bonferroni-adjusted p-value; cluster: Seurat cluster identifier; gene: gene symbol; annotation: corresponding functional annotation.

**Table S2.** Top 10 differentially expressed genes per cluster in 6hpi sample.

| p_val | avg_log2FC | pct.1 | pct.2 | p_val_adj | cluster | gene | annotation |
| --- | --- | --- | --- | --- | --- | --- | --- |
| 2.04E-35 | 3.680 | 0.264 | 0.021 | 5.64E-31 | 0 | CISH | IFN responders |
| 2.34E-82 | 3.678 | 0.530 | 0.080 | 6.48E-78 | 0 | BATF2 |  |
| 4.77E-117 | 3.564 | 0.659 | 0.105 | 1.32E-112 | 0 | AC027117.2 |  |
| 3.09E-228 | 3.310 | 0.973 | 0.456 | 8.55E-224 | 0 | ID1 |  |
| 1.98E-273 | 3.254 | 0.993 | 0.497 | 5.47E-269 | 0 | IER5L |  |
| 1.62E-50 | 3.079 | 0.424 | 0.086 | 4.47E-46 | 0 | IFITM1 |  |
| 4.47E-28 | 3.004 | 0.278 | 0.065 | 1.24E-23 | 0 | TRIM22 |  |
| 1.16E-131 | 2.887 | 0.787 | 0.269 | 3.21E-127 | 0 | ADM |  |
| 1.54E-88 | 2.789 | 0.614 | 0.146 | 4.26E-84 | 0 | FLRT3 |  |
| 2.10E-50 | 2.728 | 0.640 | 0.370 | 5.81E-46 | 0 | FGG |  |
| 1.06E-246 | 9.915 | 0.265 | 0.002 | 2.94E-242 | 1 | CCL4L2 | IFN inducers |
| 0 | 9.487 | 0.884 | 0.018 | 0 | 1 | IFNB1 |  |
| 0 | 9.413 | 0.935 | 0.059 | 0 | 1 | IFNL1 |  |
| 0 | 9.233 | 0.905 | 0.027 | 0 | 1 | IFNL2 |  |
| 0 | 9.100 | 0.899 | 0.027 | 0 | 1 | IFNL3 |  |
| 0 | 9.031 | 0.581 | 0.006 | 0 | 1 | CH25H |  |
| 0 | 8.783 | 0.781 | 0.009 | 0 | 1 | RAET1L |  |
| 0 | 8.698 | 0.933 | 0.038 | 0 | 1 | CCL5 |  |
| 0 | 8.597 | 0.406 | 0.002 | 0 | 1 | HCAR3 |  |
| 1.10E-225 | 8.262 | 0.250 | 0.003 | 3.04E-221 | 1 | TNF |  |

p_val: unadjusted p-value; avg_log2FC: average log2 fold change in expression between the indicated cluster and all remaining cells; pct.1 and pct.2: percentage of cells expressing the gene within the indicated cluster and all other cells, respectively; p_val_adj: Bonferroni-adjusted p-value; cluster: Seurat cluster identifier; gene: gene symbol; annotation: corresponding functional annotation.

**Table S3.** Top 10 differentially expressed genes per cluster in 12hpi sample.

| p_val | avg_log2FC | pct.1 | pct.2 | p_val_adj | cluster | gene | annotation |
| --- | --- | --- | --- | --- | --- | --- | --- |
| 0 | 5.284 | 0.827 | 0.141 | 0 | 0 | IFITM1 | IFN responders |
| 4.76E-188 | 5.055 | 0.399 | 0.017 | 1.31E-183 | 0 | SERPING1 |  |
| 0 | 5.037 | 0.594 | 0.035 | 0 | 0 | FLRT3 |  |
| 0 | 4.987 | 0.686 | 0.055 | 0 | 0 | AC027117.2 |  |
| 0 | 4.689 | 0.778 | 0.107 | 0 | 0 | TRIM22 |  |
| 1.00E-265 | 4.686 | 0.553 | 0.059 | 2.75E-261 | 0 | LGALS9 |  |
| 0 | 4.262 | 0.685 | 0.081 | 0 | 0 | BATF2 |  |
| 5.67E-247 | 4.117 | 0.544 | 0.070 | 1.56E-242 | 0 | MX2 |  |
| 0 | 4.096 | 0.941 | 0.299 | 0 | 0 | GDF15 |  |
| 1.96E-153 | 4.016 | 0.374 | 0.039 | 5.37E-149 | 0 | GPR37 |  |
| 5.37E-258 | 10.222 | 0.334 | 0.003 | 1.47E-253 | 1 | CCL3 | IFN inducers |
| 0 | 9.494 | 0.414 | 0.008 | 0 | 1 | CCL4L2 |  |
| 1.89E-206 | 9.427 | 0.273 | 0.002 | 5.19E-202 | 1 | CCL3L1 |  |
| 0 | 9.056 | 0.807 | 0.008 | 0 | 1 | CH25H |  |
| 0 | 8.722 | 0.954 | 0.094 | 0 | 1 | IFNL1 |  |
| 1.86E-239 | 8.674 | 0.315 | 0.003 | 5.09E-235 | 1 | KRT17 |  |
| 0 | 8.611 | 0.853 | 0.014 | 0 | 1 | HCAR2 |  |
| 0 | 8.466 | 0.997 | 0.173 | 0 | 1 | CCL5 |  |
| 0 | 8.385 | 0.817 | 0.010 | 0 | 1 | HCAR3 |  |
| 0 | 8.367 | 0.906 | 0.022 | 0 | 1 | IFNB1 |  |

p_val: unadjusted p-value; avg_log2FC: average log2 fold change in expression between the indicated cluster and all remaining cells; pct.1 and pct.2: percentage of cells expressing the gene within the indicated cluster and all other cells, respectively; p_val_adj: Bonferroni-adjusted p-value; cluster: Seurat cluster identifier; gene: gene symbol; annotation: corresponding functional annotation.

**Table S4.** Top 10 differentially expressed genes per cluster in 24hpi sample.

| p_val | avg_log2FC | pct.1 | pct.2 | p_val_adj | cluster | gene | annotation |
| --- | --- | --- | --- | --- | --- | --- | --- |
| 0 | 5.902 | 0.561 | 0.023 | 0 | 0 | SERPING1 | IFN responders |
| 0 | 5.872 | 0.814 | 0.058 | 0 | 0 | LGALS9 |  |
| 0 | 5.768 | 0.967 | 0.175 | 0 | 0 | IFITM1 |  |
| 1.46E-159 | 5.516 | 0.266 | 0.015 | 4.05E-155 | 0 | AC104035.1 |  |
| 0 | 5.286 | 0.910 | 0.115 | 0 | 0 | TRIM22 |  |
| 0 | 5.002 | 0.595 | 0.040 | 0 | 0 | MX2 |  |
| 0 | 4.683 | 0.672 | 0.148 | 0 | 0 | CP |  |
| 0 | 4.641 | 0.714 | 0.108 | 0 | 0 | CLDN2 |  |
| 0 | 4.561 | 0.873 | 0.135 | 0 | 0 | LAMP3 |  |
| 0 | 4.341 | 0.807 | 0.139 | 0 | 0 | IL18BP |  |
| 1.47E-151 | 4.240 | 0.372 | 0.078 | 4.08E-147 | 1 | IFNL2 | IFN inducers |
| 2.43E-77 | 3.937 | 0.264 | 0.077 | 6.74E-73 | 1 | IFNL3 |  |
| 0 | 3.533 | 0.760 | 0.205 | 0 | 1 | TMEM171 |  |
| 2.76E-283 | 3.359 | 0.553 | 0.097 | 7.64E-279 | 1 | HCAR2 |  |
| 0 | 3.072 | 0.807 | 0.243 | 0 | 1 | CLDN4 |  |
| 7.92E-119 | 3.042 | 0.282 | 0.052 | 2.19E-114 | 1 | NCCRP1 |  |
| 1.90E-139 | 2.944 | 0.354 | 0.078 | 5.28E-135 | 1 | CALY |  |
| 7.04E-238 | 2.855 | 0.570 | 0.141 | 1.95E-233 | 1 | HCAR3 |  |
| 7.66E-120 | 2.685 | 0.482 | 0.181 | 2.12E-115 | 1 | CXCL10 |  |
| 0 | 2.530 | 0.858 | 0.270 | 0 | 1 | FZD4 |  |
| 0 | 5.524 | 0.580 | 0.058 | 0 | 2 | CSF3 | Pro-inflammatory / Stress responses |
| 1.11E-255 | 4.742 | 0.398 | 0.030 | 3.07E-251 | 2 | BX255923.2 |  |
| 7.40E-244 | 4.613 | 0.364 | 0.024 | 2.05E-239 | 2 | BHLHE41 |  |
| 0 | 4.292 | 0.978 | 0.666 | 0 | 2 | GADD45A |  |
| 0 | 4.210 | 0.980 | 0.309 | 0 | 2 | NEURL3 |  |
| 2.03E-146 | 3.888 | 0.250 | 0.021 | 5.63E-142 | 2 | EDN2 |  |
| 0 | 3.859 | 0.507 | 0.034 | 0 | 2 | TENT5B |  |
| 2.01E-137 | 3.805 | 0.253 | 0.025 | 5.58E-133 | 2 | IL1A |  |
| 1.73E-169 | 3.787 | 0.293 | 0.026 | 4.79E-165 | 2 | UNC5B |  |
| 7.38E-275 | 3.775 | 0.444 | 0.036 | 2.04E-270 | 2 | AC140912.1 |  |
| 2.64E-134 | 3.627 | 0.571 | 0.099 | 7.32E-130 | 3 | PRPH | Cytoskeletal / Secretory |
| 1.16E-120 | 3.367 | 0.879 | 0.420 | 3.21E-116 | 3 | SNORC |  |
| 2.09E-55 | 3.227 | 0.280 | 0.050 | 5.79E-51 | 3 | DERL3 |  |
| 4.97E-131 | 3.035 | 0.826 | 0.263 | 1.38E-126 | 3 | BEX2 |  |
| 1.13E-27 | 2.762 | 0.287 | 0.095 | 3.14E-23 | 3 | SCG5 |  |
| 1.23E-128 | 2.724 | 0.954 | 0.504 | 3.42E-124 | 3 | TSC22D3 |  |
| 9.73E-37 | 2.422 | 0.294 | 0.079 | 2.70E-32 | 3 | LINC02762 |  |
| 6.73E-32 | 2.403 | 0.358 | 0.125 | 1.86E-27 | 3 | AL133476.1 |  |
| 7.49E-77 | 2.394 | 0.741 | 0.325 | 2.07E-72 | 3 | EML2-AS1 |  |
| 8.99E-64 | 2.345 | 0.543 | 0.169 | 2.49E-59 | 3 | SEPTIN5 |  |
| 0 | 8.719 | 0.305 | 0.000 | 0 | 4 | LINC00620 | Metabolic / Transporter specific |
| 0 | 8.511 | 0.524 | 0.003 | 0 | 4 | AC027088.2 |  |
| 1.37E-228 | 8.110 | 0.267 | 0.001 | 3.81E-224 | 4 | SERPINA10 |  |
| 5.04E-294 | 7.854 | 0.819 | 0.036 | 1.40E-289 | 4 | SLC22A1 |  |
| 0 | 7.573 | 0.371 | 0.002 | 0 | 4 | AL121781.1 |  |
| 6.13E-235 | 7.461 | 0.314 | 0.003 | 1.70E-230 | 4 | TYRP1 |  |
| 2.33E-219 | 7.434 | 0.276 | 0.002 | 6.46E-215 | 4 | AL138701.2 |  |
| 9.21E-132 | 7.143 | 0.305 | 0.009 | 2.55E-127 | 4 | STMN4 |  |
| 5.04E-166 | 6.640 | 0.343 | 0.009 | 1.40E-161 | 4 | AP001043.1 |  |
| 2.77E-216 | 6.585 | 0.324 | 0.004 | 7.68E-212 | 4 | AC016924.1 |  |

p_val: unadjusted p-value; avg_log2FC: average log2 fold change in expression between the indicated cluster and all remaining cells; pct.1 and pct.2: percentage of cells expressing the gene within the indicated cluster and all other cells, respectively; p_val_adj: Bonferroni-adjusted p-value; cluster: Seurat cluster identifier; gene: gene symbol; annotation: corresponding functional annotation.

**Table S5. qPCR Primer List**

| **Gene** | **Forward Primer (5'→3')** | **Reverse Primer (5'→3')** |
| --- | --- | --- |
| ***ARNTL1*** | CCAATCCATACACAGAAGCAAAC | GGTCACATCCTACGACAAACA |
| ***ARNTL2*** | GTAATGTAGTGGAGTGGGAAGAG | GCTGAGGCACAGAGTTTAAATG |
| ***CLOCK*** | CTTAGGGAACTGGCTCTGATTT | CACTGCCGTCCTGAAGTAAA |
| ***PER1*** | TCTTCCAGGATGTGGATGAAAG | TTGTGGATAGCCAGCATGAG |
| ***NR1D1*** | GTCATCCTCCTCCTCCTTCTA | TGATGTTGCTGGTGCTCTT |
| ***NR1D2*** | CAGGAGGTGTGATTGCCTATATC | GAGGAGGAGGACTGGAAACTAT |
| ***BHLHE41*** | GTTTCTCACGTTGCAACCTATTC | TCCATCTCCTTCCCTGACTT |
| ***KLF9*** | GGACATCTGTTCTTCCATACGA | CCCAAACTCCTCACTCACTAAA |
| ***NOCT*** | CTAGTACCCACCCACCTATCA | CTTCCCATTTGAGTGCTTCAAC |
| **Sendai NP** | TGCCCTGGAAGATGAGTTAG | GCCTGTTGGTTTGTGGTAAG |
| **IL29 (IFN-λ1)** | CGCCTTGGAAGAGTCACTCA | GAAGCCTCAGGTCCCAATTC |
| **RSV G** | AACATACCTGACCCAGAATC | GGTCTTGACTGTTGTAGATTGCA |
| ***OASL*** | AAATGCTCCTGCCTCAGAAA | GGGACAGAGATGGCACTGAT |
| ***TRIM22*** | CTTTATGGCTGTGCCTCCC | GTAGATGAGTGCTCCGTGGTT |
| ***CCL5*** | CCAGCAGTCGTCTTTGTCAC | CTCTGGGTTGGCACACACTT |
| ***IFNB1*** | GTCAGAGTGGAAATCCTAAG | ACAGCATCTGCTGGTTGAAG |
| ***IFI27*** | TGCTCTCACCTCATCAGCAGT | CACAACTCCTCCAATCACAACT |
| ***ISG15*** | TCCTGGTGAGGAATAACAAGGG | GTCAGCCAGAACAGGTCGTC |

qPCR primer set for circadian clock genes (*ARNTL1/2, CLOCK, PER1, NR1D1/*2, *BHLHE41, KLF9, NOCT*), innate immune/antiviral response genes (*OASL, TRIM22, CCL5, IFNB1, IFI27, ISG15, IL29*), and viral targets (SeV NP, RSV G)
